# Early warning signals are hampered by a lack of critical transitions in empirical lake data

**DOI:** 10.1101/2023.05.11.540304

**Authors:** Duncan A. O’Brien, Smita Deb, Gideon Gal, Stephen J. Thackeray, Partha S. Dutta, Shin-ichiro S. Matsuzaki, Linda May, Christopher F. Clements

**Author notes:** **Corresponding author:** Duncan A. O’Brien, **Email:**.

## Abstract

Quantifying the potential for abrupt non-linear changes in ecological communities is a key managerial goal, leading to a significant body of research aimed at identifying indicators of approaching regime shifts. Most of this work has built on the theory of bifurcations, with the assumption that critical transitions are a common feature of complex ecological systems. This has led to the development of a suite of often inaccurate early warning signals (EWSs), with more recent techniques seeking to overcome their limitations by analysing multivariate time series or applying machine learning. However, it remains unclear whether regime shifts and/or critical transitions are common occurrences in natural systems, and – if they are present – whether classic and second-generation EWS methods predict rapid community change. Here, using multitrophic data on nine lakes from around the world, we both identify the type of transition a lake is exhibiting, and the reliability of classic and second generation EWSs methods to predict whole ecosystem change. We find few instances of critical transitions in our lake dataset, with different trophic levels often expressing different forms of abrupt change. The ability to predict this change is highly technique dependant, with multivariate EWSs generally classifying correctly, classical rolling window univariate EWSs performing not better than chance, and recently developed machine learning techniques performing poorly. Our results suggest that predictive ecology should start to move away from the concept of critical transitions and develop methods suitable for predicting change in the absence of the strict bounds of bifurcation theory.

## Main

Natural ecosystems can display abrupt and non-linear shifts in state and function (Brothers *et al*. 2013; Dakos *et al*. 2015) which impairs societal activities dependent upon them. If these abrupt changes are sufficiently large to lead to local ecosystem degradation (Sguotti & Cormon 2018) and negative socio-economic impact (Crépin *et al*. 2012), then they are often classified as ‘regime shifts’ (Cooper *et al*. 2020). Concerns around the potential for regime shifts are growing as they are anticipated to increase in frequency at a global scale in response to climatic changes (Drijfhout *et al*. 2015), with the potential to cascade across systems (Rocha *et al*. 2018).

Managing ecosystems prone to these shifts is therefore vital, but doing so is challenging using current analytical tools. Resultingly, effective detection tools for characterising oncoming regime shifts are desirable, particularly cost-effective approaches accessible to all economic statuses.

Critical transitions (a.k.a catastrophic transitions) have regularly been suggested as the process driving regime shifts, via tipping points and positive feedback loops (Ludwig *et al*. 1978; Wissel 1984; Scheffer *et al*. 2001), and have led to a suite of techniques aiming to characterise oncoming transitions (Dakos *et al*. 2012a). These so-called Early Warning Signals (EWSs) are derived from bifurcation theory which states that a system can flip between two or more stable states across a critical/tipping point. EWSs attempt to detect this critical point by the phenomenon of Critical Slowing Down (CSD), which is expected to increase as a critical transition is approached (Wissel 1984). Unfortunately, the success of EWSs has been mixed despite widespread interest and research effort. For example, EWSs are consistently successful in simulated systems (Dakos *et al*. 2012b; Kéfi *et al*. 2013), whereas limited examples exist in natural settings (Biggs *et al*. 2009; Rogers *et al*. 2018; Harris *et al*. 2020). Indeed, assessments across empirical time series have been poor in their prediction of change (Burthe *et al*. 2016; Gsell *et al*. 2016), leading to doubts about the practical utility of EWSs for ecosystem managers.

Major criticisms of EWSs include their focus on ‘transitioning’ systems (Boettiger & Hastings 2012), data pre-processing (Dakos *et al*. 2012b), and, in our opinion, the assumption that critical transitions are common in natural systems (Spears *et al*. 2017). This latter is a fundamental question for the practicality of such methods in real-world data - how often do critical transitions drive regime shifts? There is a difference in definition between ‘regime shifts’ and ‘critical transitions’, but the two are often assumed the same. In its simplest form, a regime shift is ‘the process whereby an ecosystem changes from one alternative stable state to another’ (Capon *et al*. 2015) whereas critical transitions are but one possible mechanism of regime shift which can only be inferred if an incremental and slow change in a control parameter can be shown to induce a large change in state (Wissel 1984). Alternative mechanisms of regime shifts include stepwise changes in state resulting from a stepwise change in the control parameter, or noise-induced transitions that occur despite no changes in control parameter. Similarly, despite the current paradigm that lakes can exist in multiple stable states (Scheffer *et al*. 1993), neither lake chlorophyll nor global metanalysis suggest these systems consistently display multiple equilibria (Hillebrand *et al*. 2020; Davidson *et al*. 2023), a key requirement for a critical transition. Many studies assessing EWS capability in empirical data (Burthe *et al*. 2016; Gsell *et al*. 2016) may therefore be hindered as the indicators have been developed to solely quantify transition risk under certain conditions. As such, more precise classifications of lake transitions are necessary to disentangle critical transitions from these alternative regime shift mechanisms (Dakos *et al*. 2015) and allow robust assessments of EWS ability in empirical data.

An additional complication for EWS users is that, theoretically, regime shifts do not occur uniformly across all taxa (Patterson *et al*. 2021). Most regime shift or alternative stable state research focus on single representative measures of the system such as total abundance (Davidson *et al*. 2023), total cover (Hirota *et al*. 2011) or biomass (Dakos *et al*. 2012a), when multiple measures are more insightful. Recent theoretical work in multi-species systems indicate certain life-history characteristics can result in taxa being more or less illustrative of CSD (Patterson *et al*. 2021), while the interaction strength between taxa can mask transitions (Boerlijst *et al*. 2013). Reliance on univariate time series therefore makes it challenging to define an ecosystem’s dynamics as stable or transitioning, particularly as most previous work neglects non-transitioning time series (Boettiger & Hastings 2012). Consequently, although the data is available, explicit assessment of real-world critical transitions across taxa, trophic levels and transition types independently is ultimately lacking.

Fortunately, major technical developments have been made in the field which aim to circumvent these limitations. These include new computation techniques (Clements & Ozgul 2016), exploitation of information from multiple time series (Weinans *et al*. 2021) and machine learning (Bury *et al*. 2021; Deb *et al*. 2022). We therefore have access to three forms of EWS: univariate, multivariate and machine learning. In brief, univariate EWSs assess population level transitions by quantifying the presence of CSD (Dakos *et al*. 2012a), multivariate EWSs pool information from multiple time series to yield a community/system level assessment of CSD (Dakos 2018; Weinans *et al*. 2021), whereas machine learning exploit other unmeasured but learnt characteristics of the population level time series to report the probability of transition (Bury *et al*. 2021; Deb *et al*. 2022). Multivariate EWSs should logically improve the reliability of transition prediction as the maximum information and data can be generically exploited with little-to-no required knowledge of the system. However, as with classical EWSs, these developments have generally only received testing in simulated systems (Chen *et al*. 2019; Weinans *et al*. 2021) supplemented with cherry-picked empirical transition data (Deb *et al*. 2022). There is therefore a knowledge gap on how modern techniques perform in data relevant to managers and against previous methods.

In this work we examine how system state responds to changes in a control parameter (driver of upcoming regime shifts) to accurately classify regime shifts in nine long-term lake monitoring datasets (Figure 1A). We focus upon freshwater lakes as these systems are pivotal in the development of ecological theory and regime shift research (Carpenter 2003) while also providing long term and high resolution data sufficient to disambiguate trophic levels and display regime shifts. From these lake classifications, we aim to identify the ubiquity of critical transitions across lakes and planktonic trophic levels and appraise the practicality of generic EWS usage in both transitioning and non-transitioning time series. We also test novel EWS techniques’ success rates (Figure 1B,C) while optimising their potential strength via data pre-processing. The precise classification of lake fate reveals that many accepted regime shifts are not critical transitions, with different trophic levels responding uniquely to environmental change. Most EWSs therefore perform poorly, although multivariate indicators are superior to univariate.

**Figure 1.**
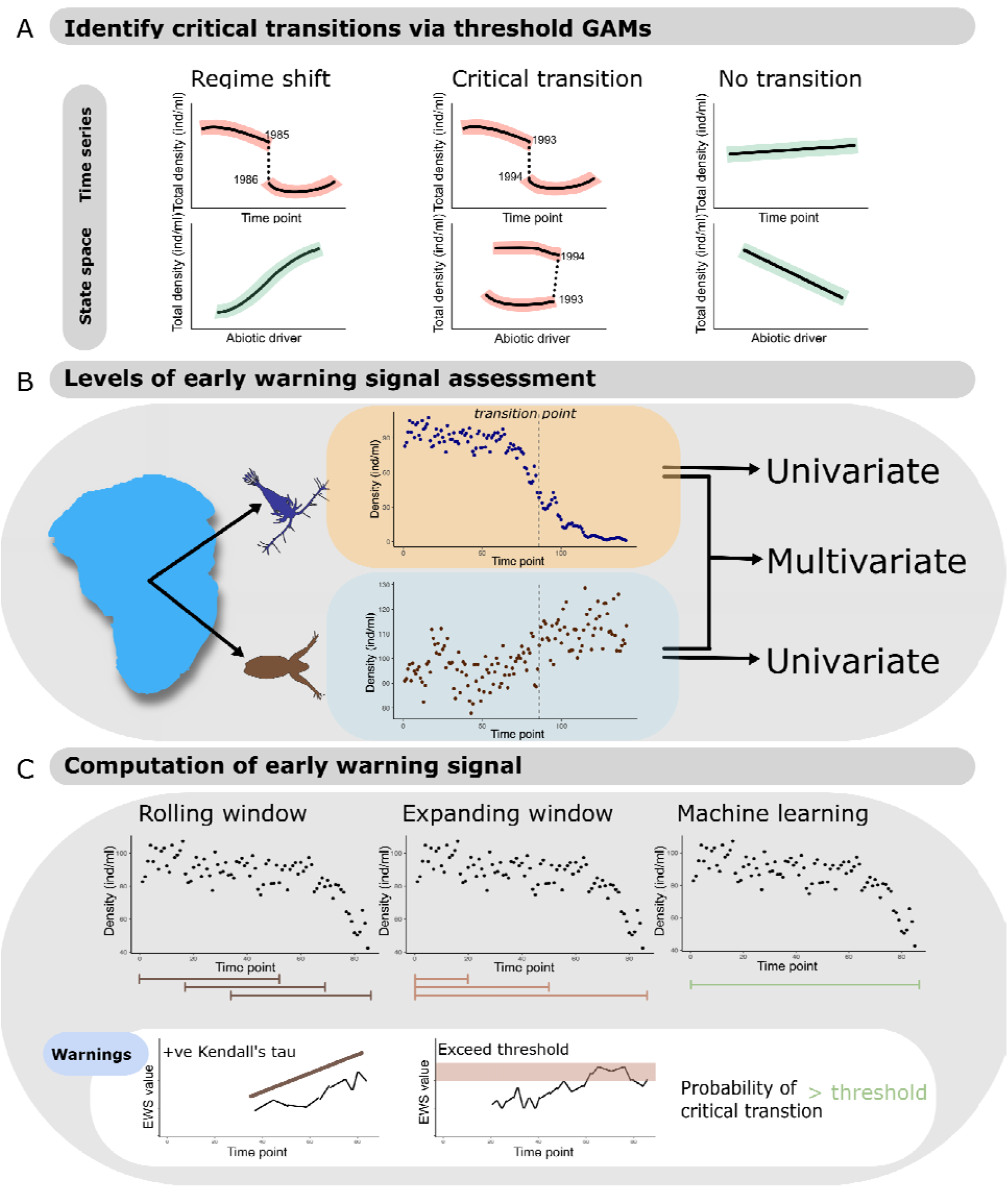
Overview of generalised additive model (GAM) and early warning signal (EWS) techniques applied in this study. A) Conceptual diagram of threshold GAM fits and how they relate to hypothesised lake fates. A regime shift can occur when there is a sudden shift in the total plankton density time series, but is not necessarily a critical transition if there is no equivalent sudden shift in the state space (i.e. in response to a small change in the abiotic driver). Only when both these criteria are fulfilled can a critical transition be hypothesised. B) Two forms of EWS can be performed: univariate EWS that consider one time series at a time and multivariate which combine information from multiple sources. The former can only give a representation of the community’s state as assessment occurs at the species level, whereas the latter assesses at the community level. C) The three computation techniques for calculating EWSs and ‘warnings’. Rolling windows are the classical form of EWS and sequentially exploit a set proportion of the time series to calculate trends in the EWS – a strong positive correlation with time indicates an oncoming transition. Expanding windows incrementally introduce new data, with the rolling average of the EWS only signalling a warning if a threshold is transgressed. And machine learning models which predict the probability of a transition based upon its knowledge of its training data. Machine learning is limited to univariate time series whereas the other computation methods can be applied to univariate or multivariate EWSs.

## Results

### Lake data

Nine publicly available and long term lake datasets were accessed - Lake Kasumigaura (Takamura & Nakagawa 2012; Takamura *et al*. 2017), Lake Kinneret (Zohary *et al*. 2014), Loch Leven (Carvalho *et al*. 2015; Gunn *et al*. 2015), Lower and Upper Lake Zurich (Pomati *et al*. 2020), Lake Mendota (Carpenter *et al*. 2017a, b), Lake Monona (Carpenter *et al*. 2017a, b), Lake Washington (Francis *et al*. 2014), and Windermere (Thackeray *et al*. 2015) – spanning a range of longitudes and environmental conditions. Planktonic data and environmental variables were extracted, standardised (see Materials and Methods), and averaged to monthly and yearly resolutions before being separated into phytoplankton and zooplankton trophic levels. Following standardisation, plankton genus richness ranged from 3 – 57 phytoplankton genus (median = 22) and 2 – 9 zooplankton genus (median = 4) in each lake.

### Quantifying lake dynamics

Using threshold generalised additive models (TGAMs), we identified optimal break points in each lake’s total phytoplankton and total zooplankton density through both time and the environmental ‘state-space’ (see Materials and Methods). This state-space was represented by the first principal component of a principal component analysis of water surface temperature, nitrate concentration and total phosphorous concentration. Namely, by cross referencing plankton densities with environment, we can identify the necessary abrupt system state response to an incremental environmental change to classify critical transitions. When comparing estimated break points between the time series and environmental models, we identified coherence in Lake Kasumigaura’s zooplankton, Lake Kinneret’s phytoplankton, Lake Monona’s zooplankton, and Lake Washington’s phytoplankton (Table S1). We consequently classified these regime shifts as critical transitions (Figure 1A). Many of the other lakes displayed breakpoints in their time series (e.g. Lake Mendota’s and Windermere’s zooplankton) but these were not matched in the environmental state space (Figure 2, Figure S1). An example analysis is provided in Figure 2 highlighting these differences. Classifications made by TGAMs were then used to ground truth downstream EWS assessments and trim time series to pre-transition data. The final time series length for each lake consequently varied between 9 – 30 years (median = 17) depending on the TGAM identified transition year, grouped into 35 transitioning time series and 209 nontransitioning time series.

**Figure 2.**
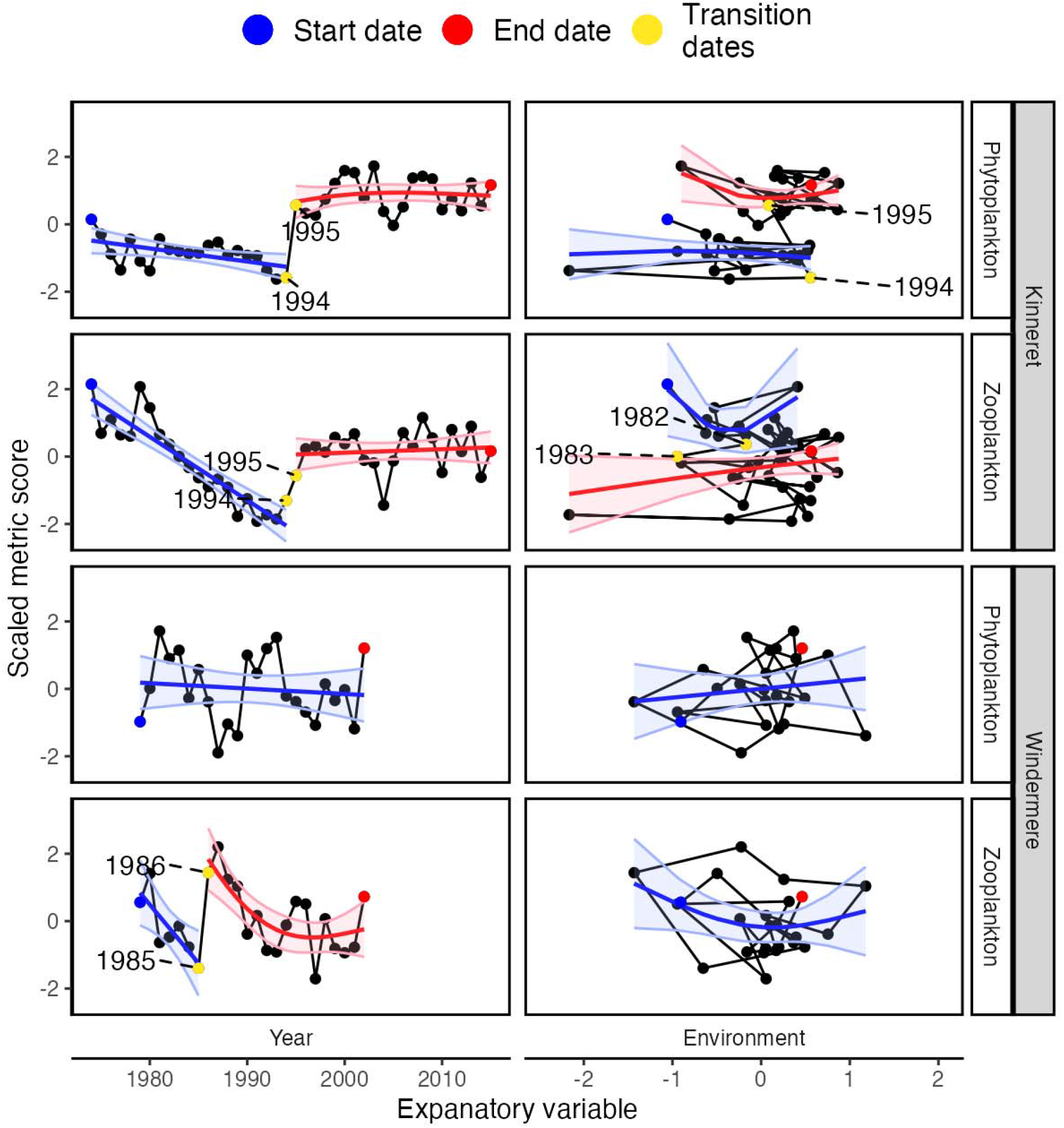
Threshold generalised additive model (TGAM) fits in Lake Kinneret and Windermere. Black points and lines represent the raw time series of plankton density in both the temporal and environmental state spaces. Start, end, and transition points are indicated by coloured points, with the dates of breakpoints also reported. Curved lines and shaded regions are the TGAM fits and 95% confidence intervals respectively. Lake Kinneret’s phytoplankton is considered the only time series displaying a critical transition due the shared break point in both the temporal and environmental model fits. The remainder of lakes can be found in Figure S1.

### Early warning signal pre-processing

Prior to EWS comparison, three detrending and deseasoning methods were performed to identify the optimal combination of detrending and deseasoning for each form of EWS. Specifically, we maximised the probability of correct prediction in the transitioning lakes, to represent the ‘best case scenario’ for EWSs. The detrending methods used included none, linear, LOESS (locally estimated scatterplot smoothing) and gaussian kernel, while deseasoning methods included none, average, time series decomposition and STL (seasonal trend estimation using LOESS). Each technique is regularly used in EWS assessments (Dakos *et al*. 2012a; Gama Dessavre *et al*. 2019) and were applied to the data factorially.

We then used a Bayesian mixed effects binomial model to test the influence of each preprocessing technique on EWS ability relative to EWS assessments made upon the raw time series. Details of the model structure can be found in the Materials and Methods. Across the five forms of EWS tested (univariate rolling window, univariate expanding window, multivariate rolling window, multivariate expanding window, and machine learning), each displayed optimal performance under different pre-processing methods (Table S3-S7). In univariate EWS, rolling windows displayed the highest prediction probabilities under linear detrending and STL deseasoning (0.08 median estimate improvement over none-none detrending and deseasoning; Table S3) whereas expanding windows performed best under no detrending and decomposition deseasoning (0.21; Table S4). Conversely, in multivariate rolling window EWSs, gaussian detrending and no deseasoning displayed the largest improvement over no pre-processing (0.21, Table S5), and gaussian-average pre-processing was optimal for multivariate expanding windows (0.24, Table S6). Finally, for machine learning models, gaussian detrending and no deseasoning improved the probability of correct classification most relative to the raw time series (0.22; Table S7). Time series that underwent these pre-processing methods were then taken forward into cross-method comparisons, assuming the best-case data scenario.

### Early warning signal overall ability

Bayesian binomial models were then fitted to test the ability of each EWS computation method and individual indicators to correctly predict the transition fate (critically transitioning versus non-transitioning) of the lake plankton time series. From this section onwards, binomial model estimates have been inverse-logit transformed from log odds to probabilities to improve interpretability. For raw model estimates please refer to Tables S7-S13 and Figures S2-S10 for model diagnostics.

When assessments are pooled across indicators, lakes, trophic levels, and resolutions, univariate EWSs estimated using expanding windows displayed the highest average probability of correct classification (Figure 3, Tables S8 and S9). This results from these EWSs displaying the highest probabilities in yearly data (median [95% credible interval] = 0.90 [0.80 – 0.95], Figure 3B) and second highest probabilities in monthly data (0.63 [0.57 – 0.68], Figure 3A). Multivariate EWSs estimated using expanding windows displayed the highest probability in monthly data (0.64 [0.56 – 0.72]) and second highest in yearly data (0.74 [0.55 – 0.86]). The remainder of computation methods were not strongly different from a 50% prediction probability although univariate rolling window EWSs were worse than chance in yearly data (0.32 [0.18 – 0.49]).

**Figure 3.**
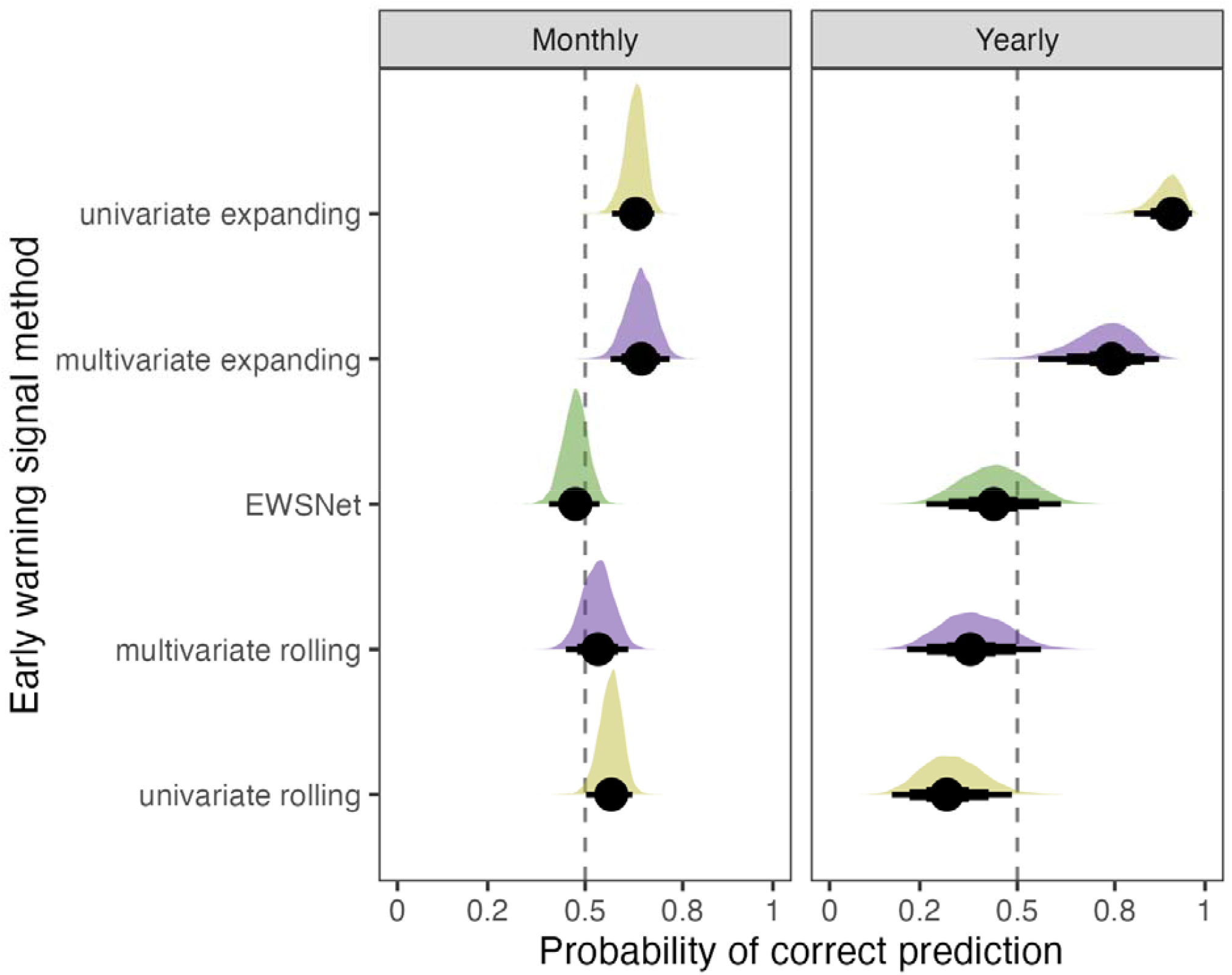
Prediction success of lake plankton fate according to early warning signal computation method displayed as density plots of posterior distributions for time series level estimates of prediction ability grouped by computation method. Computation methods are ranked by their mean ability across monthly and yearly data. The dashed vertical line shows the zero-slope - i.e. 50-50 chance of correct prediction - and each density plot represents 1000 samples from the posterior distribution of the parameter estimates. The reported values are the posterior density median values (circles), with 50% (thickest bars), 80% and 95% (thinnest bars) credible intervals back transformed from log odds to probabilities. Densities are therefore asymmetrical due to the sigmoidal relationship between log odds and probability.

Prediction probabilities were more consistent across computation methods in monthly data than yearly, with a mean prediction probability of 0.57 ± 0.07 standard deviations compared to 0.56 ±0.25. Univariate and multivariate rolling window EWSs especially declined in ability as data resolution increased. The machine learning model EWSNet was consistent across data resolutions at ∼50%.

### Individual indicator ability

Splitting these computation method level trends into indicator trends within transitioning and non-transitioning time series separately highlights how many of the above are driven by either strong true positive or true negative ability (Figure 4, Tables S8-S13). For example, unscaled EWSNet displayed a 0.93 and 0.89 true positive probability in monthly and yearly data respectively (Figure 4A), but a 0.19 and 0.05 true negative probability (Figure 4B). Scaled EWSNet displayed the inverse trend (true positive: monthly = 0.13, yearly = 0.07; true negative: monthly = 0.76, yearly = 0.97).

**Figure 4.**
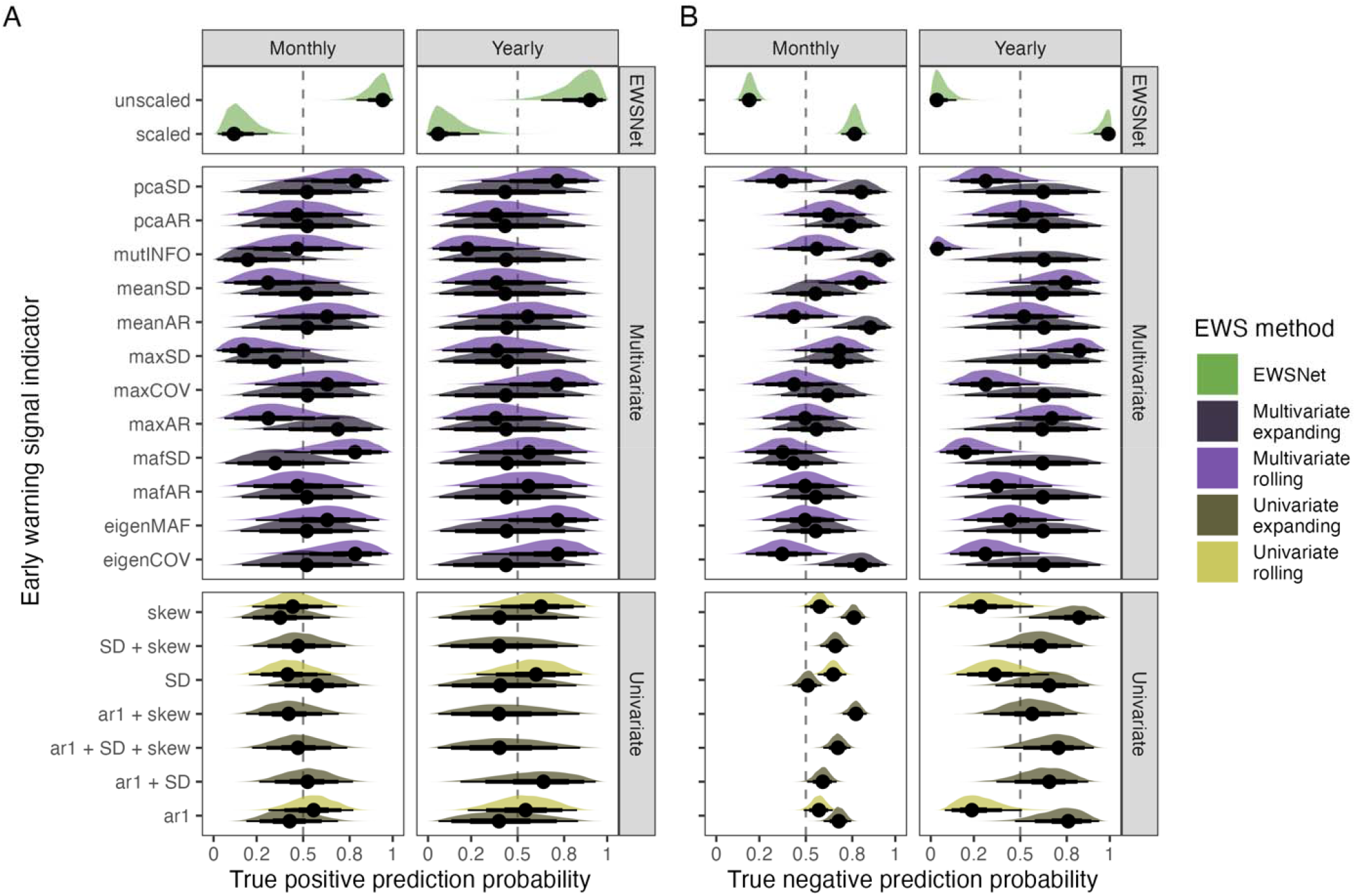
Prediction success of lake plankton fate according to individual early warning signal indicators displayed as density plots of posterior distributions for time series level estimates of prediction ability. Estimates have been segregated in to estimates made upon A) transitioning data and B) non-transitioning data. These models therefore represent indicator true positive prediction ability and true negative prediction ability respectively. The dashed vertical line shows the zero-slope - i.e. 50-50 chance of correct prediction - and each density plot represents 1000 samples from the posterior distribution of the parameter estimates. The reported values are the posterior density median values (circles), with 50% (thickest bars), 80% and 95% (thinnest bars) credible intervals back transformed from log odds to probabilities. Densities are therefore asymmetrical due to the sigmoidal relationship between log odds and probability. Colour has been used to categorise the early warning signal computation technique (i.e. rolling window, expanding window, machine learning).

To clarify, we are only focussing on the median estimates here rather than the credible intervals due to the low replication of multivariate indicators. The model is weighted by the number of trials but for multivariate EWSs, time series are concatenated to a single assessment, inflating the uncertainty of multivariate indicators relative to univariate.

There was no coherent relationship between the method of EWS calculation (rolling versus expanding windows) across variates (univariate versus multivariate). Multivariate rolling window displayed the highest mean true positive probability when averaged across all indicators in monthly time series (mean ± standard deviation: 0.54 ± 0.20) while univariate rolling displayed the highest probability in yearly time series (0.59 ± 0.04). Conversely, multivariate expanding window EWSs were superior in non-transitioning monthly time series (0.67 ± 0.15) as were univariate expanding window EWSs in non-transitioning yearly time series (0.68 ± 0.09).

This therefore suggests that individual indicators are highly variable and so should be considered individually. However, the method of calculation alters the indicators’ sensitivity. For example, mutual information was consistently poor in its predictions, displaying a, <0.5 median probability in all circumstances excluding in the monthly non-transitioning time series (expanding window = 0.90, rolling window = 0.56). Composite univariate EWSs (Drake & Griffen 2010) computed via expanding windows (e.g. ar1 + SD, ar1 + SD + skew) were relatively reliable across resolutions and transition types but non-composite indicators struggled in transitioning yearly data. Multivariate rolling window indicators such as max covariance (maxCOV), PCA variance (pcaSD) and the dominant eigenvalue of the maximum autocorrelation factor dimension reduction (eigenMAF) were particularly effective in transitioning time series but weak in non-transitioning data, whereas only the expanding window indicators maximum and mean autocorrelation (maxAR, meanAR) were consistent across transitioning and non-transitioning data.

## Discussion

Motivated by debates surrounding multiple stable states in ecology and the need for reliable and generic critical transition detection tools, we assessed the prevalence of critical transitions in a range of empirical lake systems. We then compared the ability of current state-of-the-art early warning signal (EWS) methods to correctly predict ecosystem fate regardless of transition type or trophic level. We found that multiple regime shifts were identifiable across our lake network, but only a proportion of these were critical transitions. Similarly, within a transitioning lake, phytoplankton and zooplankton trophic levels could display different transition mechanisms; one may critically transition whereas the other experiences a step change/transient. Together, these findings may have hampered previous attempts to test EWS ability. Multivariate EWS methods outperformed univariate, and the method of data pre-processing also dramatically influenced their ability. Machine learning models performed variably depending on their data processing although had extremely high true positive rates. Overall, computational and methodological advances have improved EWS ability in empirical data over classical approaches but still are not sufficiently reliable for generic usage due to conceptual concerns regarding the ubiquity of appropriate systems.

Generic EWS predictive ability is limited as it appears some system specific knowledge is necessary. Namely, understanding the potential for a critical transition/multiple stable states and identifying a mechanistic driver of transition are particularly key. These are not new arguments (Boettiger & Hastings 2012; Dakos *et al*. 2015) but are worth reiterating to avoid the conflation of critical transitions with any form of abrupt change/regime shift. The threshold generalised additive model approach used here does identify the same regime shifts as previous studies (Roelke *et al*. 2007; Francis *et al*. 2014; Fukushima & Arai 2015; Walsh *et al*. 2017) but only some of these were classifiable as critical transitions. We suggest it is prudent to consider that temporal dynamics of driver variables (e.g. nutrient concentration) are themselves non-linear (Manzoni *et al*. 2004) and display pulse events (Collins *et al*. 2008). For example, an anomalous year may push a system away from equilibrium in to a long transient, as potentially occurred during 1985 in Windermere’s zooplankton (Figure 2), or a step change in environmental conditions can result in novel communities (Conley *et al*. 2007). Consequently, critical transitions can occur earlier, later, or not at all, even if a regime shift occurs.

Bifurcation theory similarly assumes that a system exists in a parameter space where multiple states/attractors are present (Wissel 1984). The conflicting evidence for the presence of multiple stable states (Roelke *et al*. 2007; Capon *et al*. 2015; Hillebrand *et al*. 2020; Davidson *et al*. 2023) challenges the validity of this assumption in ecological data and any bifurcation detection techniques. Indeed, long transients may represent a superior conceptual model for lake regime shifts (Morozov *et al*. 2020) due to the non-stationary dynamics of driving variables (Manzoni *et al*. 2004; Collins *et al*. 2008), and recent identification of rate induced tipping (Ashwin *et al*. 2012) implies that a system can exhibit a sudden regime shift once the driver rapidly varies above a critical rate. R-tipping states that a rapid change in the driver parameter can cause a sudden shift even without crossing a bifurcation point. It is associated with the movement of a quasi-static attractor and a critical rate of change. There is consequently no ‘one-size-fits-all’ approach to regime shift pre-emption and focus should shift to either representing the stability landscape (Dakos *et al*. 2015; Ushio *et al*. 2018) and/or describing a system mechanistically (Boerlijst *et al*. 2013).

These concerns are compounded by our finding that different components of the system (trophic levels here) can display distinct transition forms. Such incoherence is not unexpected, with certain taxa being more informative than others (Boerlijst *et al*. 2013; Patterson *et al*. 2021; Medeiros *et al*. 2022) but is rarely considered. Monitoring one representative variable is therefore unlikely to be sufficient and many facets of the system should be tracked, including plausible environmental drivers. In lake systems this is established practice but is more difficult to apply to other ecological biomes/scenarios. Drawing upon autonomous monitoring will contribute to this gap (Besson *et al*. 2022), but requires implementation.

Our EWS findings broadly agree with previous attempts to assess EWSs in aquatic systems (Burthe *et al*. 2016; Gsell *et al*. 2016) although those studies solely focus on univariate rolling window indicators. Indeed, our approach of disentangling critical transitions from regime shifts and inclusion of non-transitioning systems strengthens conclusions that classic EWSs are inappropriate in transitioning time series. We do however show that these EWSs often correctly classify non-transitions, a point not previously clarified due to an almost exclusive focus in transitioning data (Boettiger & Hastings 2012). Conversely, the unexpected ability of expanding window univariate EWSs in yearly data similarly supports the recommendations of Clements *et al*. (Clements & Ozgul 2016) for using composite signals when data quality is poor. This approach is successful as multiple EWS indicators are combined and the EWS’s sensitivity can be tailored to match a manager’s conservatism to transition risk. Univariate signals therefore do have limited value but are best suited to situations where high resolution data is not available, and variability is low. Practical examples include threatened animal populations (Clements *et al*. 2017) and vegetation patterning changes in response to drought (Eby *et al*. 2017).

In contrast to univariate EWSs, multivariate signals appear the most reliable way of characterising critical transitions, balancing the ratio of true positive to true negative predictions. This is not unreasonable as pooling information increases the likelihood of encountering taxa displaying critical slowing down (CSD) - though see Pilotto *et al*. (Pilotto *et al*. 2020) for arguments that broad scale or global pooling weakens the identifiability of biodiversity trends. We circumnavigate these concerns by focussing our multivariate measures on lake level data, so that any interpretation is at the local level where critical transitions are likely to occur. However, requiring a broad coverage of the system of interest is necessary..

Machine learning has also been touted as having high potential for classifying critical transitions (Bury *et al*. 2021; Deb *et al*. 2022; O’Brien *et al*. 2022), by identifying empirical phenomena distinct from CSD. We focussed on the EWSNet model due to its coverage of multiple transition types and mathematical models, and accessibility in a number of different coding languages (O’Brien *et al*. 2022), but the approach performed poorly in our empirical plankton time series. The training data provided to EWSNet is certainly a contributing factor as evidenced by the differences between the two forms of EWSNet tested here; a strong true positive but poor true negative success rate in the default unscaled model was reversed in the scaled form. EWSNet’s authors (Deb *et al*. 2022) do advocate the finetuning of machine learning models with data of similar magnitude/dynamics as the target system to mitigate these issues. Here, we applied EWSNet generically without specific finetuning due to a desire to test its generic ability, plus challenges in generating appropriate training data for each lake system. This may explain the model’s low classification success. Expansion to models trained upon multivariate data may also be a solution, given the relative strength of multivariate EWSs relative to univariate. That being said, if failure to detect a critical transition is more harmful than presumptuously intervening, then EWSNet in its generic form does have merit. We do overall, however, reaffirm previous suggestions that machine learning requires tailoring to the specific system of interest and is inappropriate for generic usage. This begs the question whether if the system requires targeted modelling to finetune these signals, whether EWSs or machine learning can provide greater insight than such a model alone.

The combination of apparent EWS uncertainty and uncommonness of critical transitions strengthens the opinion that resilience measures are a superior generic tool than CSD based indicators. While some of the measures tested here are considered resilience indicators (e.g mutual information (Weinans *et al*. 2021)), more complicated measures have recently emerged independent from the assumption for local stability. For example, Ushio *et al*. (Ushio *et al*. 2018) exploit empirical dynamic modelling to estimate the Jacobian matrix of a multivariate community and extract a stability index which accurately diagnoses vulnerable periods in fish communities. This has been developed further by Medeiros *et al*. (Medeiros *et al*. 2022) to identify key species for management based upon their contribution to the system’s Jacobian. Similarly, Fisher Information (a measure of information inferred from observed data on a latent variable) appears to characterise both regime shifts (Spanbauer *et al*. 2014) and overall resilience (O’Brien *et al*. 2023). Resilience indicators have the additional benefit of not requiring post hoc detection of regime shifts nor critical transitions. Thus, practical real-time monitoring is possible regardless of critical transition risk as any unexpected loss of stability is a concern for ecosystem managers. EWSs can provide some information on resilience loss (Dakos *et al*. 2015; Weinans *et al*. 2021) but are conceptually linked to the presence of CSD (Hillebrand *et al*. 2020). Critical transitions should only conceptually occur when there are more than one state to transition across (Dakos *et al*. 2015; Hillebrand *et al*. 2020), and so EWSs should only be relevant in those circumstances.

Resilience indicators are ultimately capable of quantifying system stability both in proximity to critical transitions and in systems where transitions are not possible (Yang *et al*. 2019), lending them greater generic power than EWSs. Our findings support the conceptual use of these measures, as we show that multivariate EWSs display the highest prediction probabilities across transitioning and non-transitioning lakes.

To conclude, successfully applying and interpreting EWSs in real time is complicated and potentially unintuitive. Modern techniques struggle to be consistent and generic in empirical systems due each system’s unique dynamics and lack of techniques for identifying true real-world critical transitions. Ecosystems can display regime shifts via many forms (Dakos *et al*. 2015), the majority of which are theoretically not going to be detectable by EWSs. It therefore appears tailoring machine learning models to one’s system of interest or focussing on resilience measures (which quantify stability rather than transitions) will be more useful for ecosystem managers. In systems where critical transitions are possible, combining information is critical to maximise EWS robustness but can still fail. The interpretation of EWSs generically applied across systems therefore requires caution.

## Methods

### Lake plankton data

Weekly plankton densities and abiotic variables were sourced from nine publicly available longterm monitoring datasets: Lake Kasumigaura (Takamura & Nakagawa 2012; Takamura *et al*. 2017), Lake Kinneret (Zohary 2004), Loch Leven (Carvalho *et al*. 2015; Gunn *et al*. 2015), Lower and Upper Lake Zurich (Pomati *et al*. 2020), Lake Mendota (Carpenter *et al*. 2017a, b), Lake Monona (Carpenter *et al*. 2017a, b), Lake Washington (Francis *et al*. 2014), and Windermere (Thackeray *et al*. 2015). As these datasets encompassed a range of institutions, we performed a standardization procedure. Unidentified and/or unnamed species were removed and if a species was not recorded on a sampling date, that species’ density was assumed to be zero. The data was then averaged to mean density per month and per year. Species were then pooled to genus level to minimise the likelihood of zero densities, but a genus would be further dropped if they disappeared for a period longer than 12 months. This is necessary as downstream deseasoning will spuriously introduce cycles of non-zero densities into periods of zeroes. The final dataset for each lake consequently consisted of genus level densities across two trophic levels (phytoplankton and zooplankton) and three plausible abiotic drivers – water surface temperature (°C), nitrate concentration (μgL), and total phosphorous concentration (μgL).

Prior to early warning signal (EWS) assessment, each plankton time series was pre-processed via detrending and deseasoning using a range of techniques. Detrending is considered important for improving the reliability of EWS assessments (Gama Dessavre *et al*. 2019), so we applied three commonly used methods (linear detrending, LOESS smoothing, and gaussian smoothing) and compared assessments made to those based upon the raw time series. Linear detrending fits a linear model between time and plankton density, with the residuals of this model representing the detrended time series (Lenton *et al*. 2012). LOESS, or local polynomial regression smoothing, subtracts a smooth curve fitted by local polynomial regression of span 0.5 from the raw time series (Bury *et al*. 2021), while gaussian kernel smoothing applies a linear filter, by subtracting the weighted moving average from the raw time series (Lenton *et al*. 2012). Additionally, each time series was deseasoned as monthly plankton data is inherently seasonal (Maberly *et al*.

2022), and the repeated non-linear cycles can hinder EWS capability (O’Brien & Clements 2022). We therefore applied three deseasoning techniques (averaging, additive decomposition, and STL) factorially with the detrending methods to identify the optimal combination. Averaging simply subtracts the average value for a given month from the current data point of that month (O’Brien *et al*. 2023), additive decomposition estimates the seasonal cycle from moving averages which is then subtracted from the raw time series (Zarnowitz & Ozyildirim 2006), and stl (seasonal trend estimation using loess) which also estimates the average seasonal cycle but uses local polynomials rather than linear/moving averages (Cleveland *et al*. 1990). All data pre-processing was performed using the *EWSmethods* package (O’Brien *et al*. 2022) implemented in R 4.2.1 (R Core Team 2022).

### Critical transition pre-classification

For a sudden non-linear change in plankton density to be truly a critical transition, the system must display a sudden change in state following a small incremental change in the stressing variable (Wissel 1984). Consequently, to identify historic critical transitions, assign system ‘fates’, and allow us to assess the ability of each EWS to classify transitions, we fitted threshold generalised additive models (TGAMs) to raw, yearly total plankton densities through both time and the environmental ‘state-space’ (Figure1A, Figure 2). Together, these models allow us to disentangle critical transitions from pulse events or step changes. TGAMs fitted between time and plankton densities reveal sudden state changes, but the environmental model attempts to represent the environmental conditions the system is experiencing at each time point.

Specifically, we represented these environmental conditions as the first principal component of the lake’s abiotic drivers. We then used these TGAMs as a double validation tool where a breakpoint in the density time series indicates a non-linear step change in the system, whereas a breakpoint in the environmental state-space indicates a large change in response to a small change in stressor (Figure 1A). Therefore, we consider that a shared breakpoint is required between both the time series and state-space GAM fits for a critical transition to be identified. We also performed our classification procedure independently across phytoplankton and zooplankton trophic levels as this dataset gives us the opportunity to question whether critical transitions are shared across both components of the system. For example, it is plausible that a critical transition in one trophic level will not necessarily be matched by a critical transition in the other if the former trophic level is a driver of the second; a critical transition in one system component drives a step change in another.

Practically, we followed the TGAM fitting procedure of Ciannelli *et al*. (Ciannelli *et al*. 2004), where a breakpoint can be introduced in to a GAM smooth. The optimal location of that break is identified by minimising the generalised cross-validation (GCV) score of the model (analogously to Akaike’s information criterion), with the optimal choice between GAM and TGAM also selected via minimising GCV. To minimise the likelihood of overfitting, the total number of knots for each thin-plate spline smooth was restricted to a maximum of six, and, in the threshold form of the model, both halves of the smooth were restricted to three knots each. A threshold model was only fit if the continuous GAM smooth displayed an effective degree of freedom >= three. This represents an approximately cubic shape which plausible contains a step change and minimises the likelihood of erroneously accepting TGAMs for approximately linear models. The optimal number of knots was selected via restricted maximum likelihood, and, in both the time series and environmental models, breaks were only permitted between adjacent time points. All models were fitted through *mgcv* 1.8-40 (Wood 2004).

### Early warning signal assessments

Each plankton time series was then subjected to EWS assessments using each of the univariate, multivariate, and machine learning techniques provided in the *EWSmethods* R package (O’Brien *et al*. 2022). Table S2 details the specifics of each indicator, but overall, univariate EWSs accept individual time series and quantifying the degree of critical slowing down (CSD) for that one taxa. Multivariate EWSs expand these assessments from single time series to multiple by either averaging across univariate EWSs or by extracting CSD information from a dimension reduction of the system (Figure 1B). Finally, machine learning involves a convolutional neural network being trained upon mathematical models that undergo critical and non-critical transitions, to learn characteristics of a critical transition and report the probability of transition.

Univariate and multivariate EWSs were also assessed using two different computation approaches: rolling and expanding windows (Figure 1C). Rolling windows portion the time series into a fixed window length, which then ‘rolls’ along the time series, incrementally calculating the indicators of Table S2. The Kendall tau correlation of these indicators then represents the quantity of interest, where a ‘strong’ correlation coefficient is indicative of an oncoming transition. In this study, we maintained a rolling window size 50% the total length of the time series following Dakos *et al*. (Dakos *et al*. 2012a). The alternative expanding window computation incrementally introduces new data after a set burn in period. Each indicator is standardised by subtracting its running mean from its calculated value at time *t* before division by its running standard deviation (Clements & Ozgul 2016). A composite metric can then be constructed by summing all individual indicator values calculated per *t*. An oncoming transition is consequently identified when the indicator/composite metric exceeded its expanding mean by a certain threshold value. Here, we set that threshold at two standard deviations due its favourable performance relative to alternative threshold values (Clements *et al*. 2017). We also imposed a burn in period 50% the total length of the time series to mitigate spurious signals that occur at the beginning of assessment resulting from few data points in the window.

We also included the machine learning model EWSNet (Deb *et al*. 2022). EWSNet utilises the entirety of the pre-transition time series to provide probabilities of the likelihood of 1) a critical transition, 2) a smooth transition, or 3) no transition. We used the full ensemble of 25 models provided by EWSNet and averaged across them to improve the robustness of probability estimates following the suggestions of O’Brien *et al*. (O’Brien *et al*. 2022). We additionally tested the effect of scaled (between 1 and 2) versus unscaled data processing on the quality of EWSNet predictions.

To enable comparability between transitioning and non-transitioning taxa, lakes containing transitions were subset prior to the year identified by TGAMs. Non-transitioning lakes were subset to 85% of their total length. This ensures we can infer the near future of the non-transitioning lake correctly.

Additionally, as the various EWS method classes all generate different outputs, we converted these outputs into the binary presence-absence of a ‘warning’ (Figure 1C). For rolling window computations, a warning was accepted if a positive Kendall tau correlation was in the 95th quartile of Kendall tau correlations from a dataset permuted from the original time series (Dakos *et al*. 2012a), for expanding windows when the two standard deviation threshold was exceeded for two or more time points (Clements *et al*. 2019), and for EWSNet, when the model predicted a critical transition (i.e. the strongest probability)(O’Brien *et al*. 2022). This presence-absence of a warning was then compared to the ground-truth labels identified by the TGAMs, resulting in a binomial dataset of successes and failures.

### Early warning signal ability

To estimate the classification ability of each EWS method, we developed a series of Bayesian hierarchical models using success/failure as response variable. Early warning signal method class and the specific EWS indicator itself were explored as categorical fixed effects in separate models. Early warning signal method class was treated as a factor with five levels: univariate rolling, univariate expanding, univariate machine learning, multivariate rolling, and multivariate expanding. Indicator was also treated as a factor with 21 levels for each indicator detailed in Table S2. To account for the non-independence of repeated measurements for each lake and fate within each lake (transitioning versus non-transitioning trophic levels), we included a nested random effect of fate within lake identity.

We also tested which combination of detrending and deseasoning methods maximised the probability of a correct prediction for each indicator in the transitioning lakes. This was represented as a factor with 16 levels for each pre-processing combination. We then used this information to fit the above models using time series which underwent the optimal combination.

The resulting general model structure was:

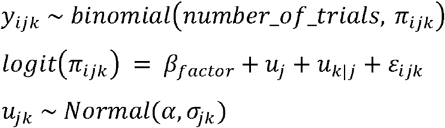

where *y*_*ijk*_ represents the log odds of correctly classifying a system’s fate in the *i*th time series, in the *j*th lake with the known fate *k*. π therefore represents the probability of the classification, β represents slopes, μ the random intercepts, and ε the remaining error. Each model was fitted without a global intercept to allow us to interpret the absolute effect of each factor level on the probability of correctly classifying a time series’ fate.

We set the weakly informative priors:

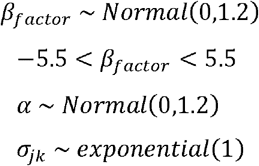

where *factor* represents the varying factors tested and *j* and *k* are lake identity and lake fate (the presence/absence of a critical transition identified by TGAM analysis) respectively. β_*factor*_ is constrained between -5.5 and 5.5 as this represents a ∼1% to ∼99% probability.

Models were written in the Stan language (Stan Development Team 2022) and implemented in *brms* 2.18.0 (Bürkner 2017), where they were run for 10,000 iterations following a 2000 iteration warmup period. Convergence was assessed via the identification of well mixed trace plots (Figures S2-S7) and appropriate Rhat values (equal to 1, Tables S3-S13), with posterior predictive checks validating the final model posterior shape relative to the observed data (Figures S8-S10). During interpretation, we back-transformed the log odds into probabilities of correct classification, and used the overlap of the posterior distribution’s credible intervals against 50% to identify EWS approaches that provide better estimates than chance. Modelling the probability of correct classification in this way rather than using true positive:false postive ratios allows us to control for other confounding factors in the dataset, namely lake identity and the varying number of trials (i.e. EWS assessments) between lakes and EWS methods.

## Acknowledgments

DAO received funding from the GW4+ FRESH Centre for Doctoral Training in Freshwater Biosciences and Sustainability (NE/R011524/1) and SD received funding from the Ministry of Education, Government of India (Prime Minister’s Research Fellowship). We thank Tamar Zohary for the long-term phytoplankton dataset from Lake Kinneret, Francesco Pomati for data from the Zurich lakes, the North Temperate Lakes LTER group for data from North American lakes, and Heidrun Feuchtmayr for Windermere data; monitoring at Windermere is currently supported by Natural Environment Research Council award number NE/R016429/1 as part of the UK-SCAPE program delivering National Capability. We also thank the field and laboratory teams who have collected all of the data used in this study.

## Notes

### Competing Interest Statement

The authors have declared no competing interest.

