## Supplementary Information for "Early warning signals are hampered by a lack of critical transitions in empirical lake data"

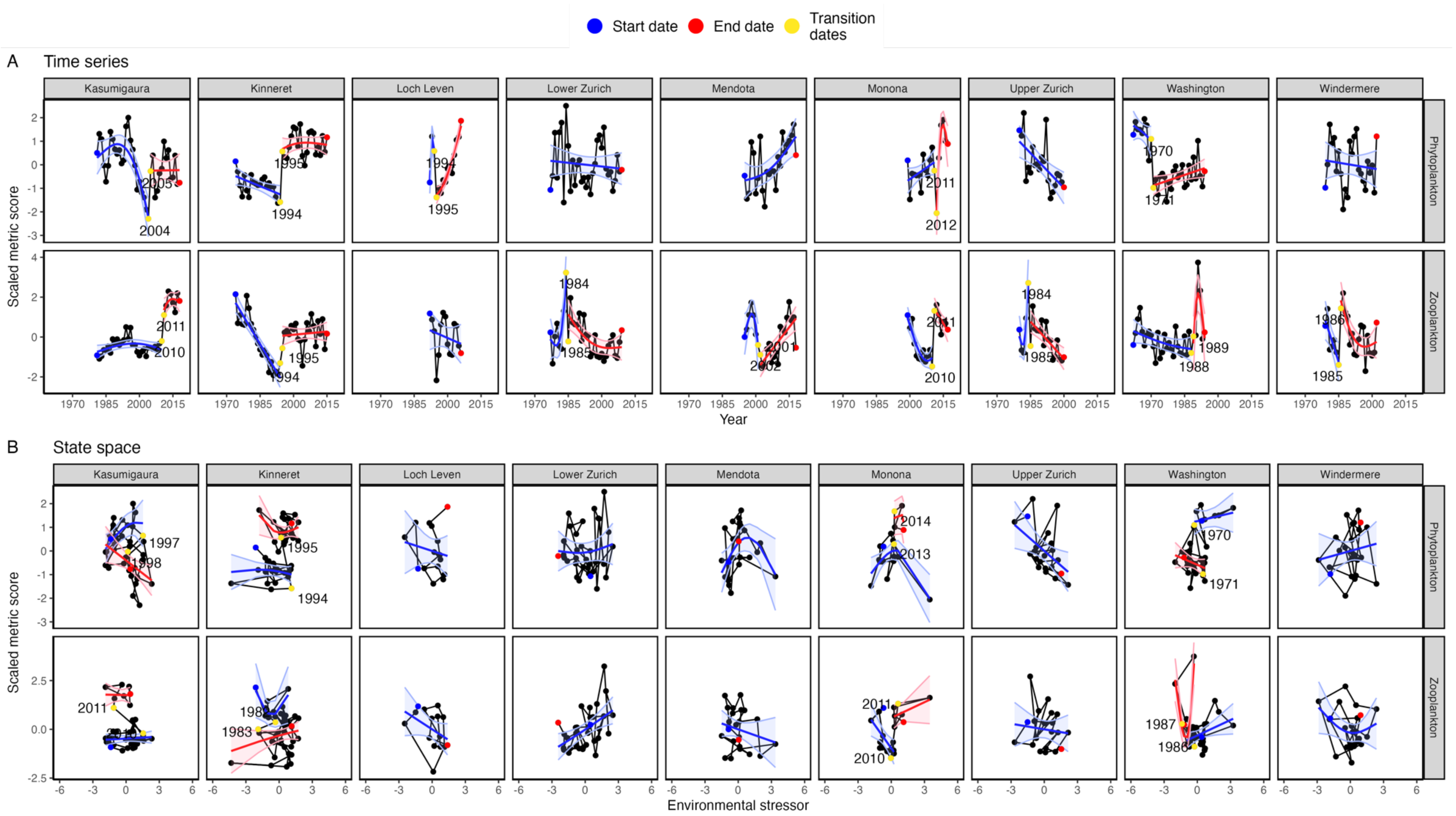


**Fig. S1.** All threshold generalized additive model fits across lakes and trophic levels. Black points and lines represent the raw time series of plankton density in both the temporal and environmental state spaces. Start, end, and transition points are indicated by coloured points, with the dates of breakpoints also reported. Curved lines and shaded regions are the TGAM fits and 95% confidence intervals respectively.

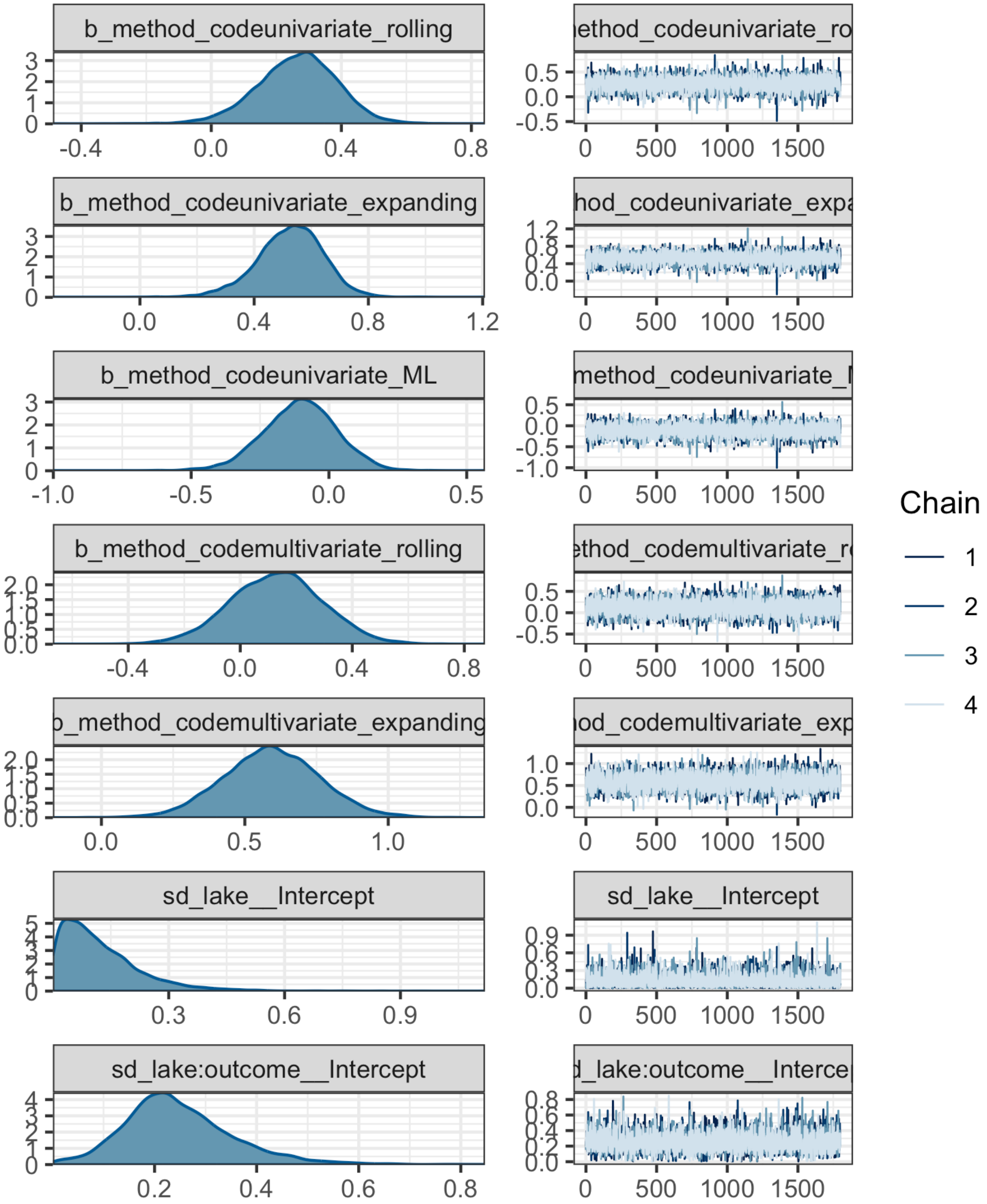


**Fig. S2.** Trace plots for each parameter of a hierarchical binomial Bayesian model fitted between early warning signal computation method and successful prediction of lake fate in monthly plankton data. Visually, a converged fit is indicated by unimodal density plots (left) and 'well-mixed'/highly overlapping chains (right).

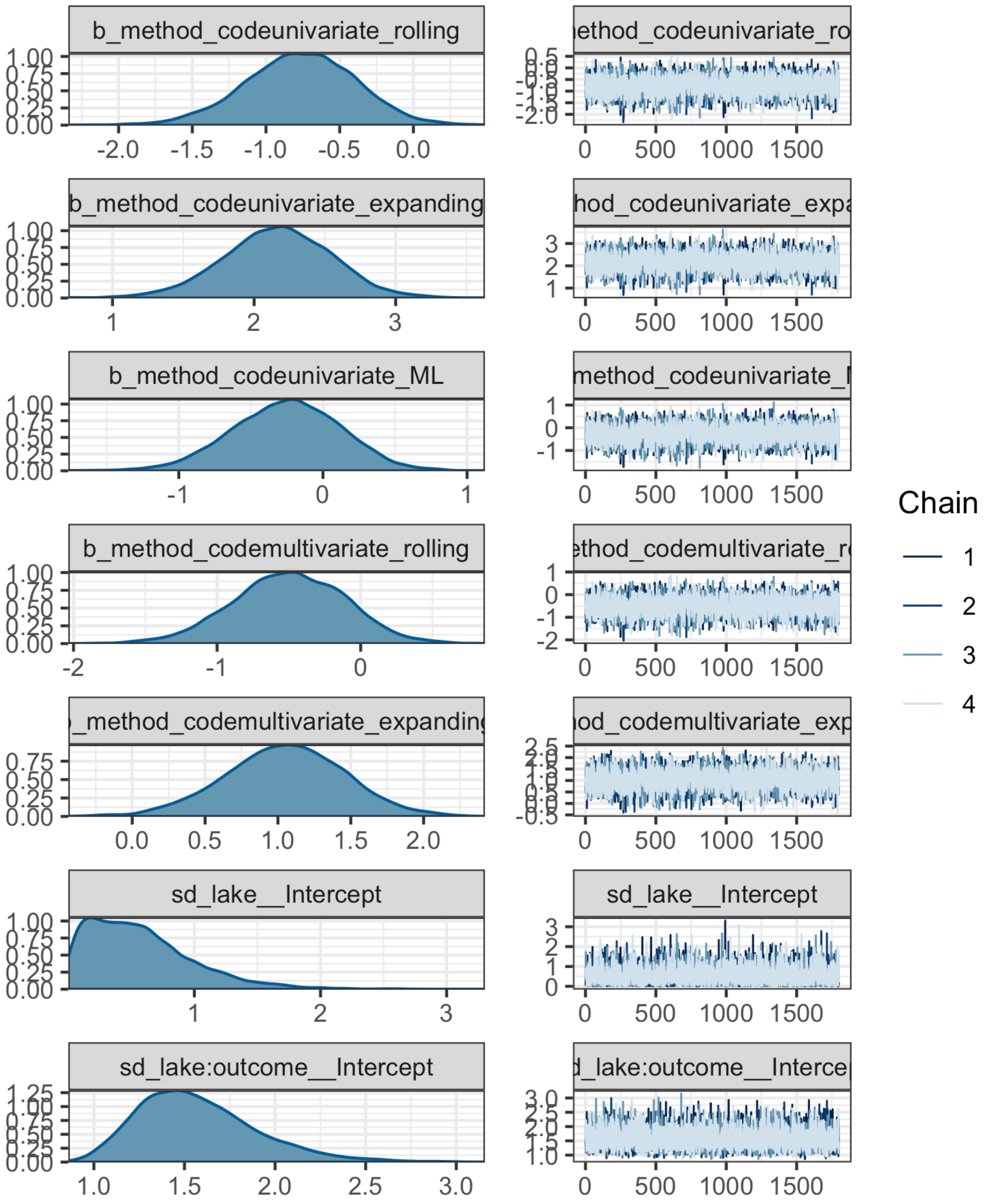


**Fig. S3.** Trace plots for each parameter of a hierarchical binomial Bayesian model fitted between early warning signal computation method and successful prediction of lake fate in yearly plankton data. Visually, a converged fit is indicated by unimodal density plots (left) and ‘well-mixed’/highly overlapping chains (right).

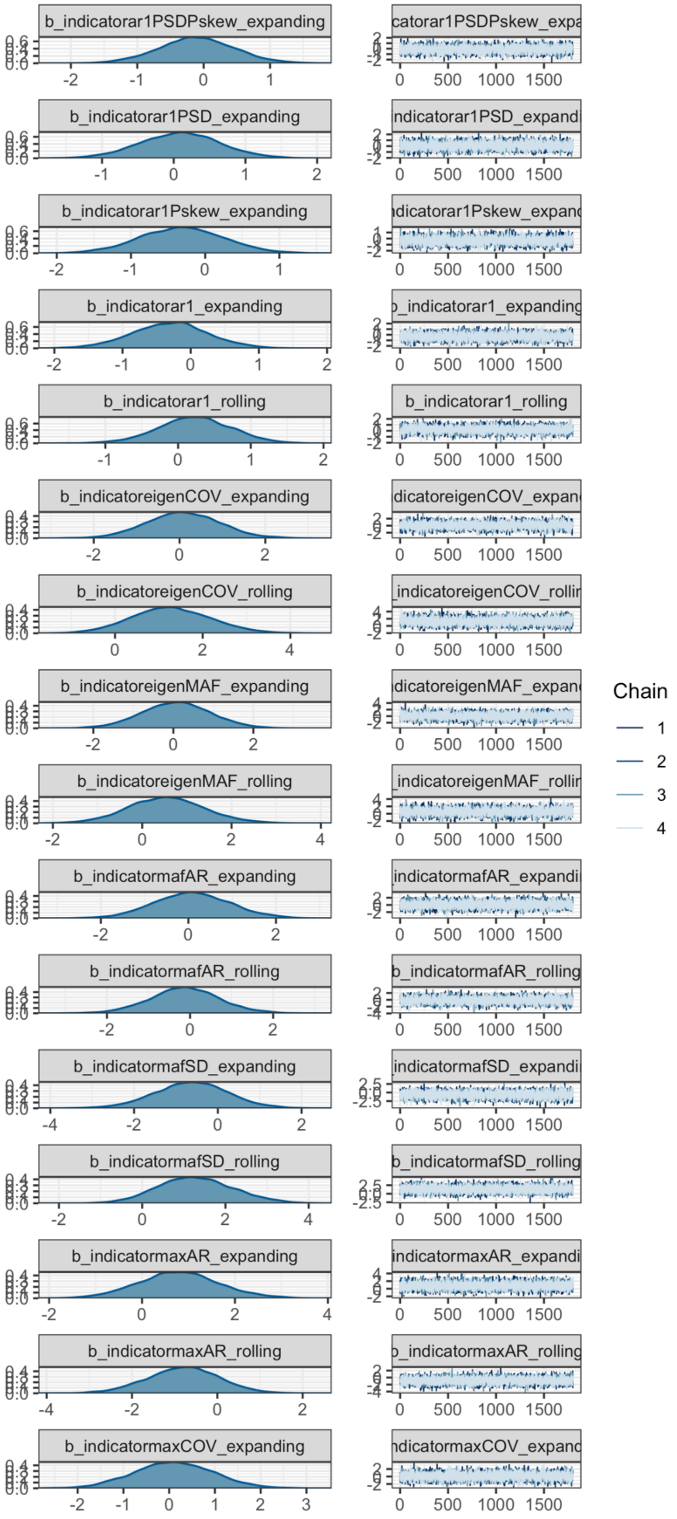


**Fig. S4.** Trace plots for each parameter of a hierarchical binomial Bayesian model fitted between early warning signal indicator and successful prediction of lake fate in transitioning monthly plankton data.

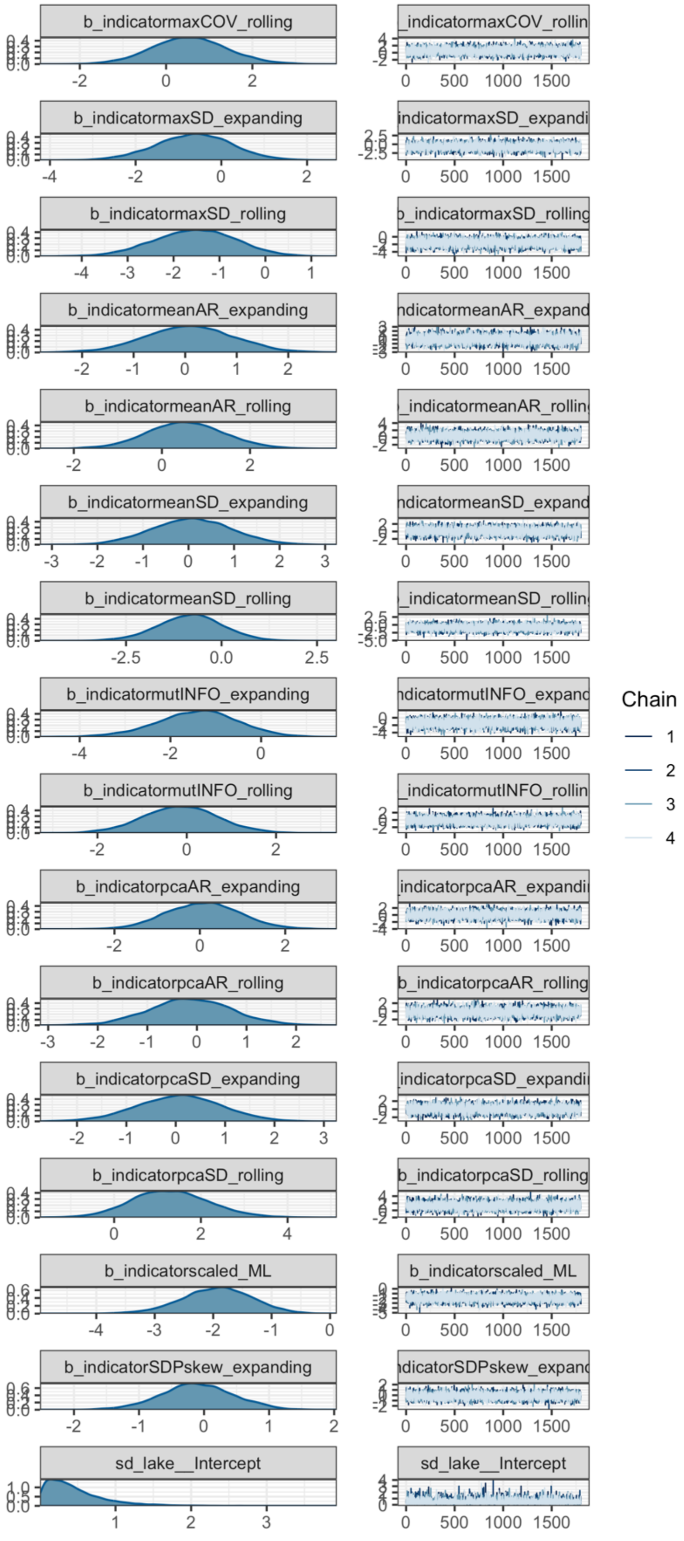
b_indicatormaxCOV_rolling
b_indicatormaxSD_expanding
b_indicatormaxSD_rolling
b_indicatormeanAR_expanding
b_indicatormeanAR_rolling
b_indicatormeanSD_expanding
b_indicatormeanSD_rolling
b_indicatormutINFO_expanding
b_indicatormutINFO_rolling
b_indicatorpcaAR_expanding
b_indicatorpcaAR_rolling
b_indicatorpcaSD_expanding
b_indicatorpcaSD_rolling
b_indicatorscaled_ML
b_indicatorSDPskew_expanding
sd_lake__Intercept
Chain
1
2
3
4

**Fig. S4 cont.**

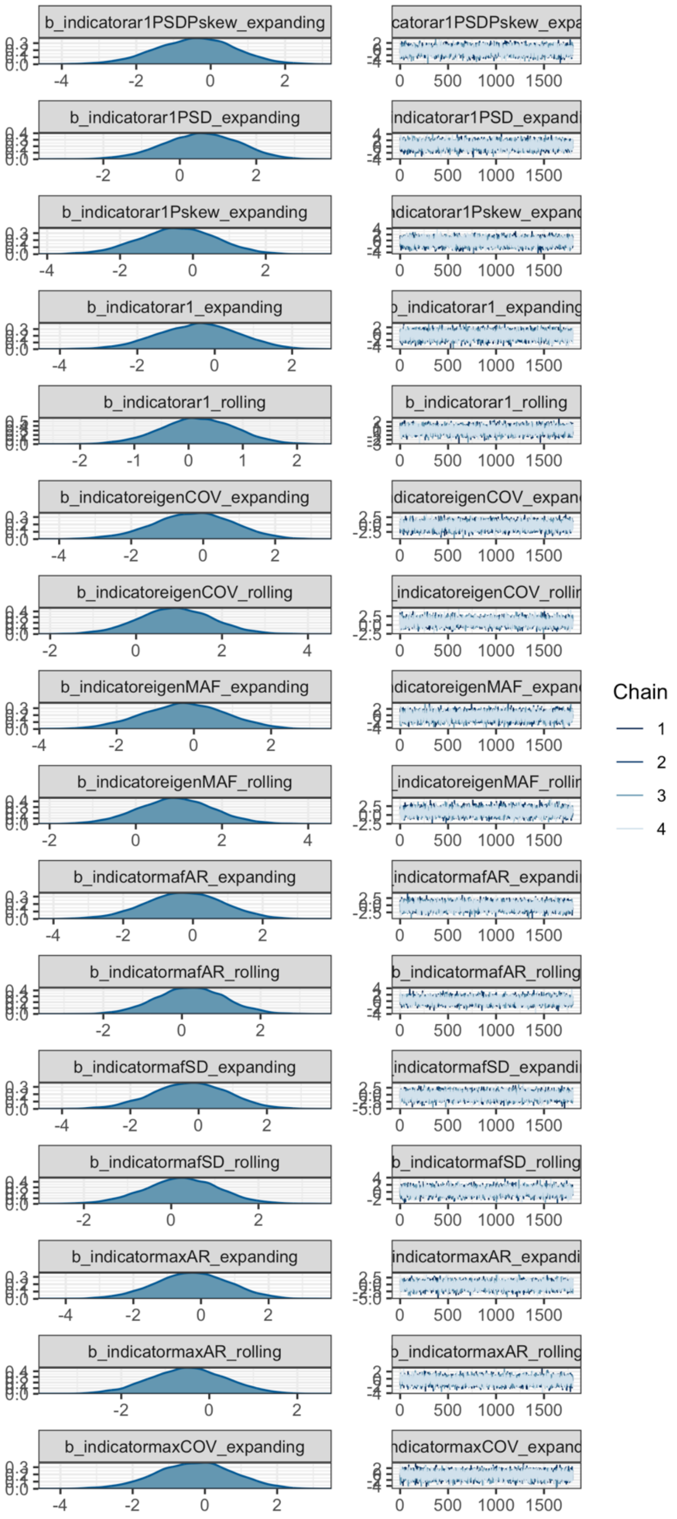


**Fig. S5.** Trace plots for each parameter of a hierarchical binomial Bayesian model fitted between early warning signal indicator and successful prediction of lake fate in transitioning yearly plankton data.

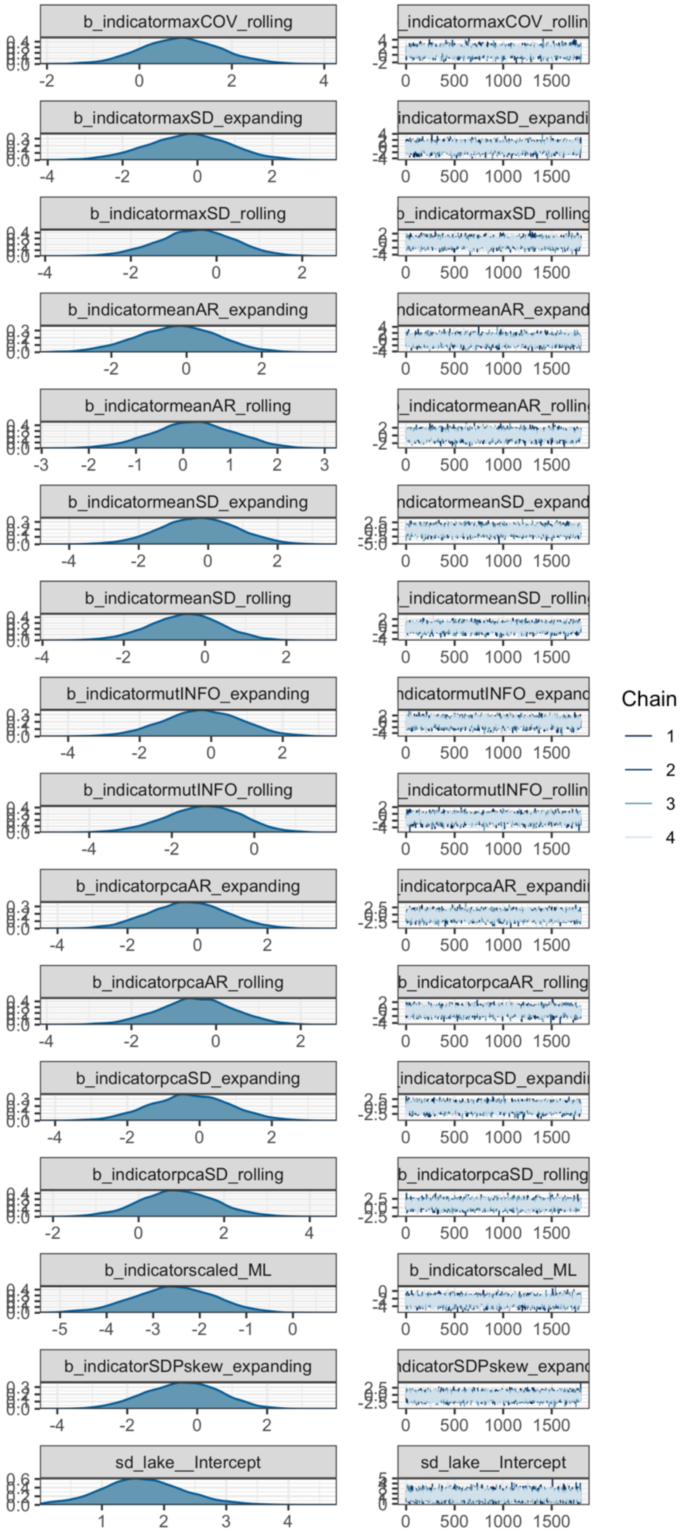
b_indicatormaxCOV_rolling
b_indicatormaxSD_expanding
b_indicatormaxSD_rolling
b_indicatormeanAR_expanding
b_indicatormeanAR_rolling
b_indicatormeanSD_expanding
b_indicatormeanSD_rolling
b_indicatormutINFO_expanding
b_indicatormutINFO_rolling
b_indicatorpcaAR_expanding
b_indicatorpcaAR_rolling
b_indicatorpcaSD_expanding
b_indicatorpcaSD_rolling
b_indicatorscaled_ML
b_indicatorSDPskew_expanding
sd_lake__Intercept
0 500 1000 1500
Chain
1
2
3
4

**Fig. S5 cont.**

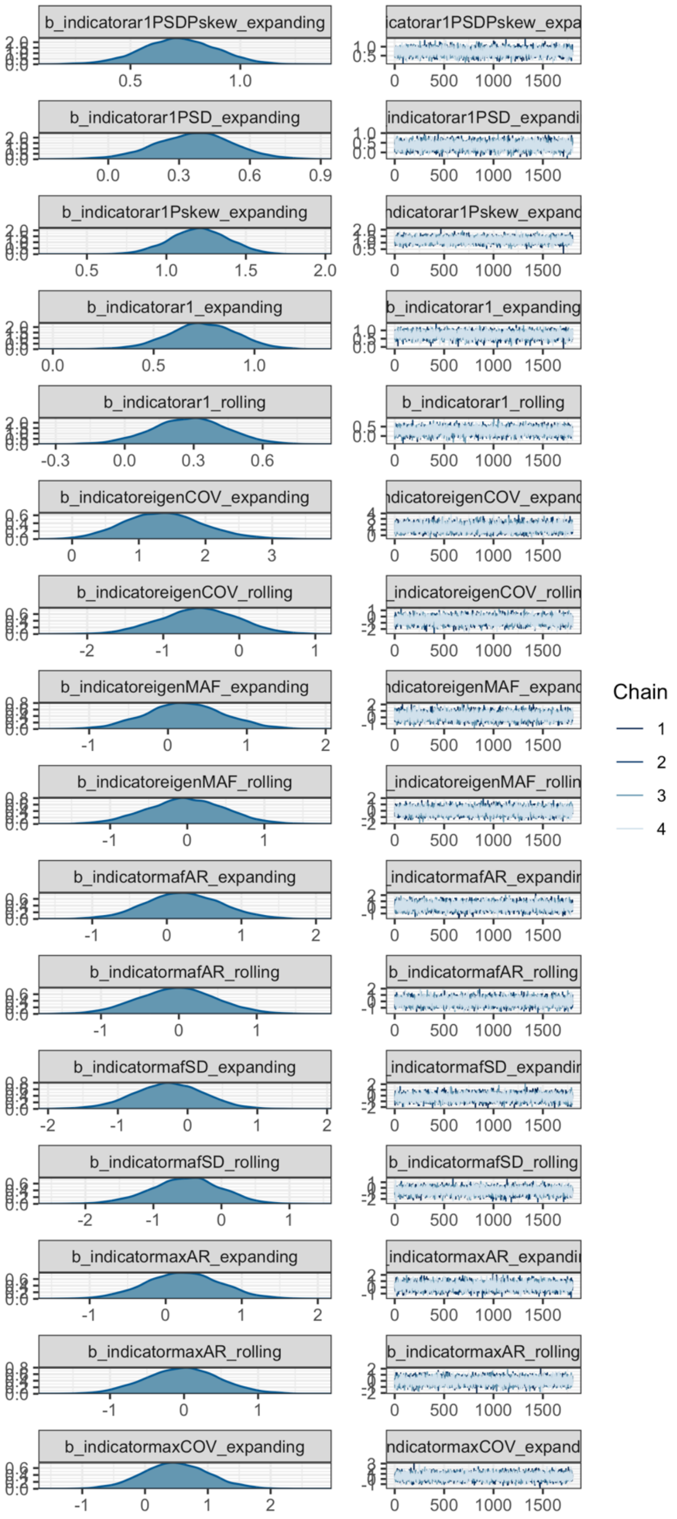


**Fig. S6.** Trace plots for each parameter of a hierarchical binomial Bayesian model fitted between early warning signal indicator and successful prediction of lake fate in non-transitioning monthly plankton data.

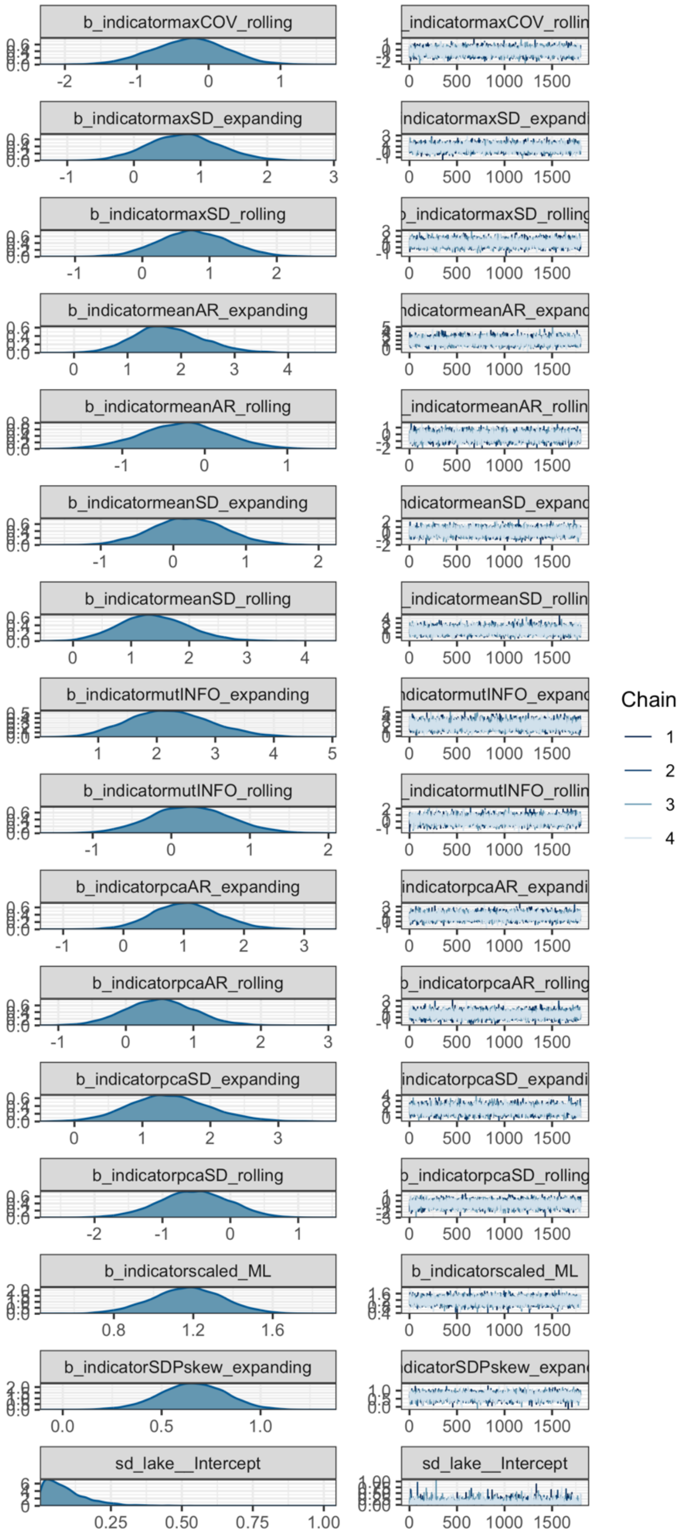
b_indicatormaxCOV_rolling
b_indicatormaxSD_expanding
b_indicatormaxSD_rolling
b_indicatormeanAR_expanding
b_indicatormeanAR_rolling
b_indicatormeanSD_expanding
b_indicatormeanSD_rolling
b_indicatormutINFO_expanding
b_indicatormutINFO_rolling
b_indicatorpcaAR_expanding
b_indicatorpcaAR_rolling
b_indicatorpcaSD_expanding
b_indicatorpcaSD_rolling
b_indicatorscaled_ML
b_indicatorSDPskew_expanding
sd_lake__Intercept
Chain
1
2
3
4

**Fig. S6 cont.**

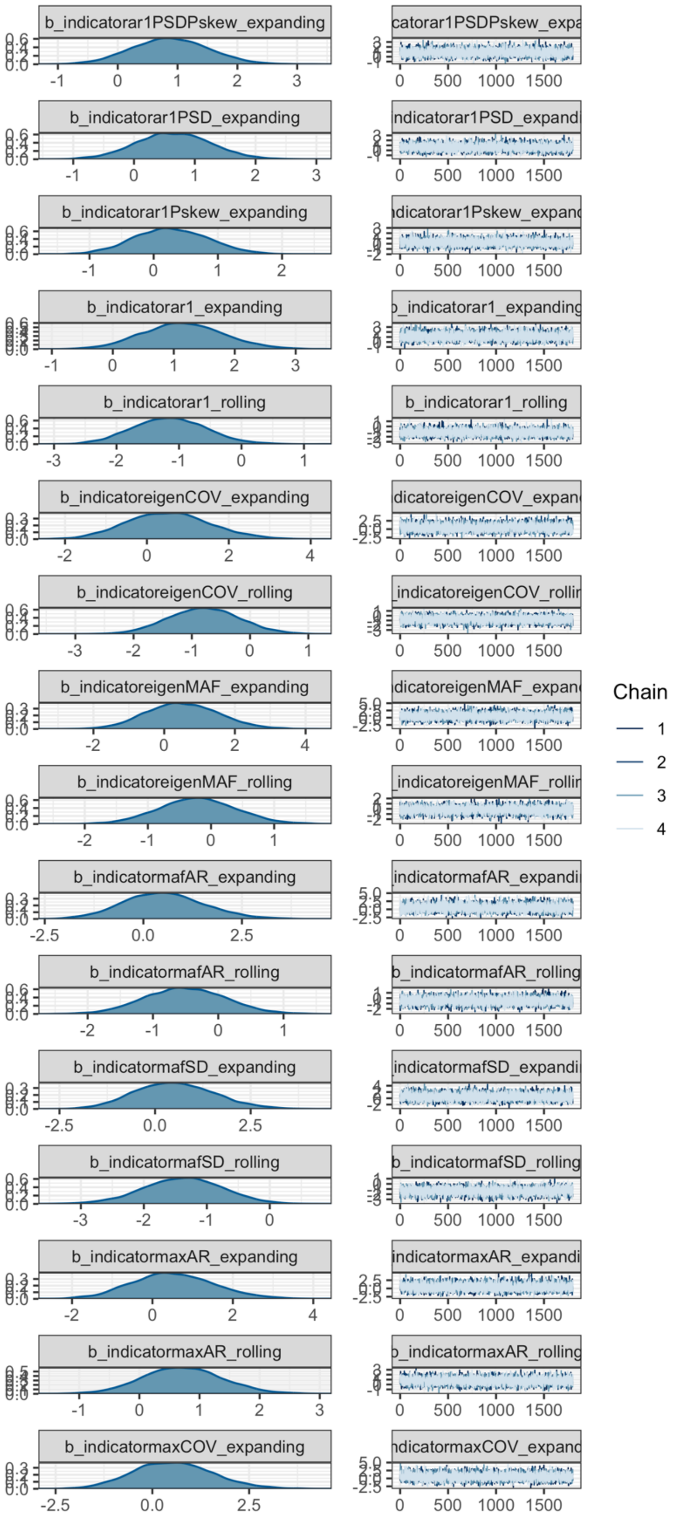


**Fig. S7.** Trace plots for each parameter of a hierarchical binomial Bayesian model fitted between early warning signal indicator and successful prediction of lake fate in non-transitioning yearly plankton data.

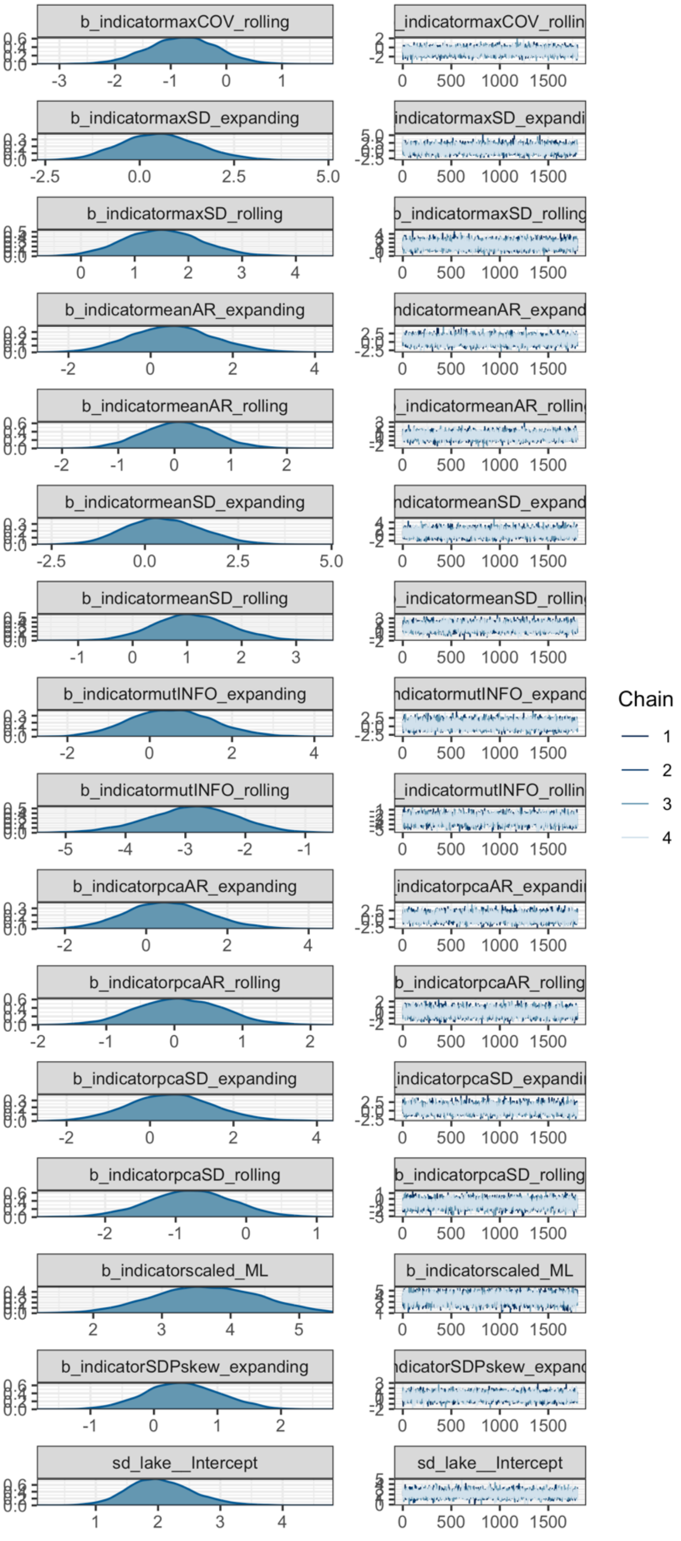
b_indicatormaxCOV_rolling
_indicatormaxCOV_rollin
b_indicatormaxSD_expanding
indicatormaxSD_expandi
b_indicatormaxSD_rolling
_indicatormaxSD_rolling
b_indicatormeanAR_expanding
ndicatormeanAR_expand
b_indicatormeanAR_rolling
_indicatormeanAR_rollin
b_indicatormeanSD_expanding
ndicatormeanSD_expand
b_indicatormeanSD_rolling
_indicatormeanSD_rollin
b_indicatormutINFO_expanding
ndicatormutINFO_expanc
b_indicatormutINFO_rolling
_indicatormutINFO_rollin
b_indicatorpcaAR_expanding
indicatorpcaAR_expandi
b_indicatorpcaAR_rolling
b_indicatorpcaAR_rolling
b_indicatorpcaSD_expanding
indicatorpcaSD_expandi
b_indicatorpcaSD_rolling
b_indicatorpcaSD_rolling
b_indicatorscaled_ML
b_indicatorscaled_ML
b_indicatorSDPskew_expanding
ndicatorSDPskew_expanc
sd_lake__Intercept
sd_lake__Intercept
0 500 1000 1500
Chain
1
2
3
4

**Fig. S7 cont.**

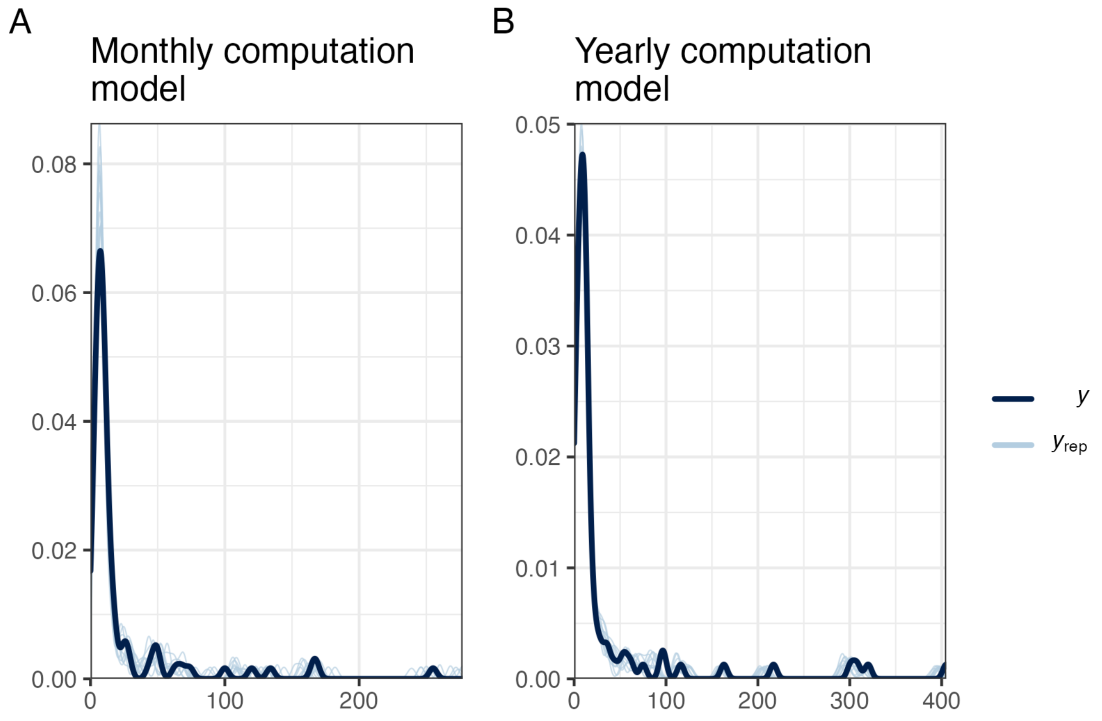


**Fig. S8.** Posterior predictive checks of hierarchical binomial Bayesian models fitted between early warning signal computation method and successful prediction in A) monthly and B) yearly plankton data. An appropriate fit occurs when *yrep* reasonably reflects *y*.

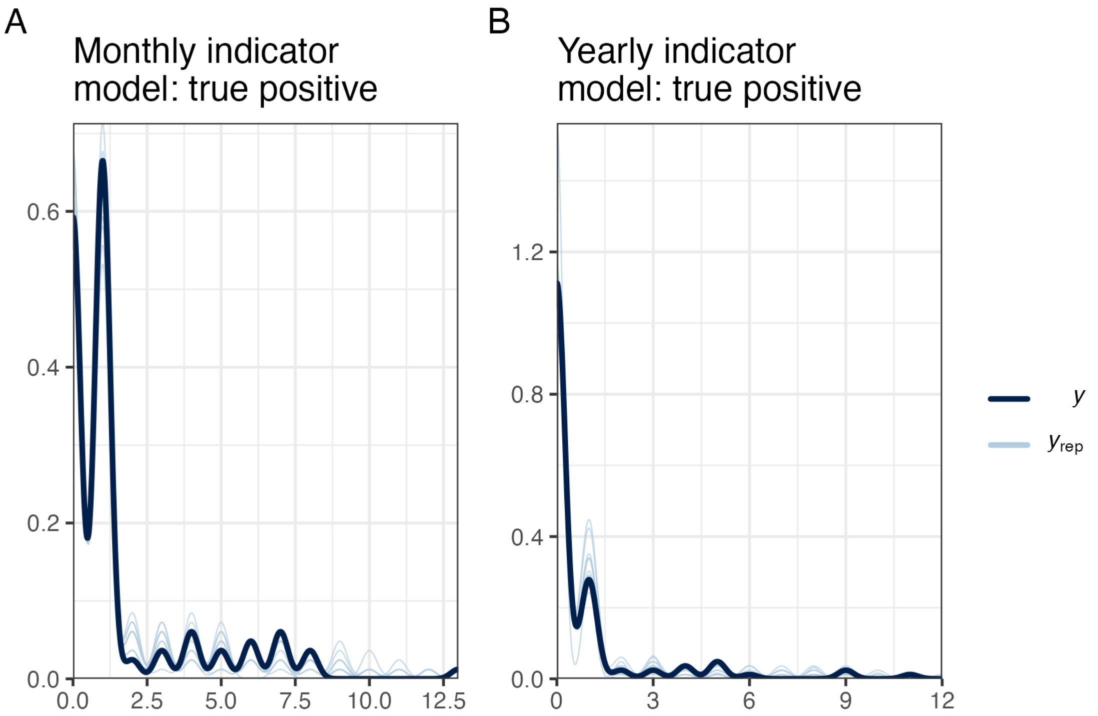


**Fig. S9.** Posterior predictive checks of hierarchical binomial Bayesian models fitted between early warning signal indicator and successful prediction in A) monthly and B) yearly transitioning plankton data. These models represent the true positive ability of each indicator. An appropriate fit occurs when *yrep* reasonably reflects *y*.

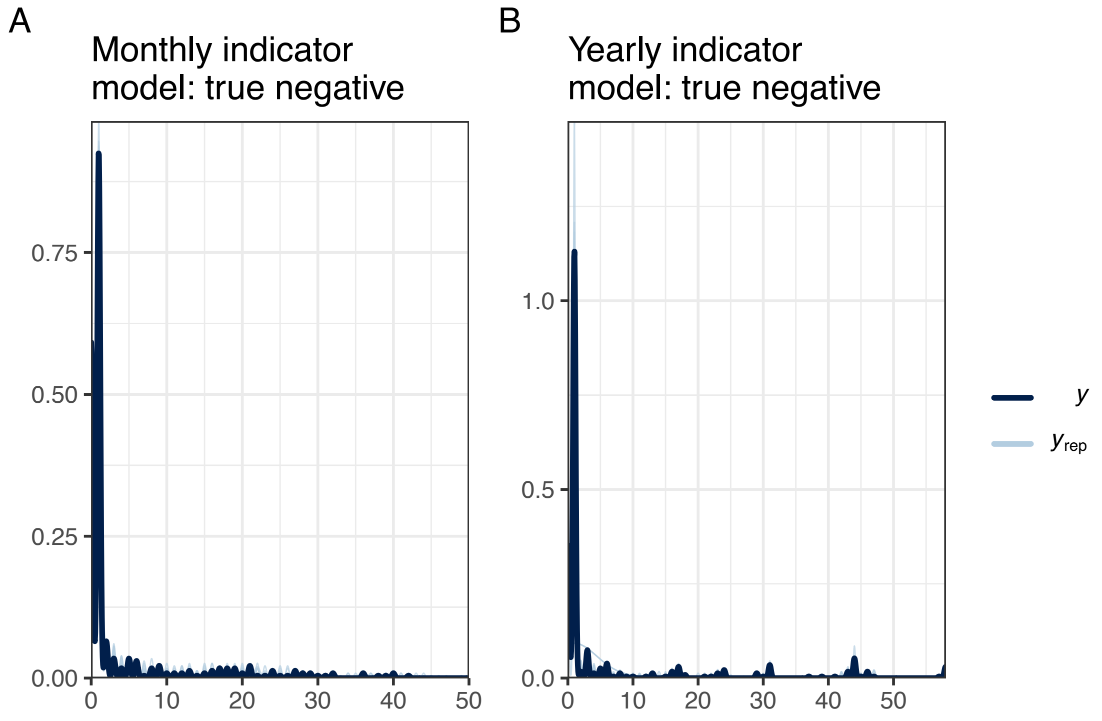


**Fig. S10.** Posterior predictive checks of hierarchical binomial Bayesian models fitted between early warning signal indicator and successful prediction in A) monthly and B) yearly non-transitioning plankton data. These models represent the true negative ability of each indicator. An appropriate fit occurs when *yrep* reasonably reflects *y*.

**Table S1. Breakpoints and critical transitions identified from threshold generalised additive models.**

| Lake | Trophic level | Explanatory variable | Breakpoint year | Critical transition identified |
|---|---|---|---|---|
| Kasumigaura | Phytoplankton | Time | 2004 | No |
| | | Environment | 1997 | |
| | Zooplankton | Time | **2010** | **Yes** |
| | | Environment | **2010** | |
| Kinneret | Phytoplankton | Time | **1994** | **Yes** |
| | | Environment | **1994** | |
| | Zooplankton | Time | 1982 | No |
| | | Environment | 1994 | |
| Loch Leven | Phytoplankton | Time | 1994 | No |
| | | Environment | NA | |
| | Zooplankton | Time | NA | No |
| | | Environment | NA | |
| Lower Zurich | Phytoplankton | Time | NA | No |
| | | Environment | NA | |
| | Zooplankton | Time | 1984 | No |
| | | Environment | NA | |
| Mendota | Phytoplankton | Time | NA | No |
| | | Environment | NA | |
| | Zooplankton | Time | 2001 | No |
| | | Environment | NA | |
| Monona | Phytoplankton | Time | 2011 | No |
| | | Environment | 2013 | |
| | Zooplankton | Time | **2010** | **Yes** |
| | | Environment | **2010** | |
| Upper Zurich | Phytoplankton | Time | NA | No |
| | | Environment | NA | |
| | Zooplankton | Time | 1984 | No |
| | | Environment | NA | |
| Washington | Phytoplankton | Time | **1970** | **Yes** |
| | | Environment | **1970** | |
| | Zooplankton | Time | 1988 | No |
| | | Environment | 1986 | |
| Windermere | Phytoplankton | Time | NA | No |
| | | Environment | NA | |
| | Zooplankton | Time | 1985 | No |
| | | Environment | NA | |

**Table S2. Description of each individual early warning signal indicator and which variate category it belongs to.**

| Early warning signal method | Indicator | Description | Abbreviation |
|---|---|---|---|
| Univariate | autocorrelation at lag-1 | The similarity between temporally adjacent data points – i.e. the correlation between the time series and the lagged version of itself. | ar1 |
| | variance | A measure of the degree of dispersion displayed in the time series. Is represented as the standard deviation here. | SD |
| | skewness | The degree of asymmetry in the distribution of values displayed in the time series. | skew |
| | composite of each combination of the above | In the expanding window computation, the three above indicators can be standardised and combined to improve their reliability. Two and three indicator combinations were performed in this study. | ar1 + SD, ar1 + skew, SD + skew, ar1 + SD + skew |
| Multivariate | mean autocorrelation at lag-1 | Average autocorrelation across all time series representing the system. | meanAR |
| | max autocorrelation at lag-1 | The strongest autocorrelation of all time series representing the system. | maxAR |
| | mean variance | Average standard deviation across all time series representing the system. | meanSD |
| | max variance | The largest standard deviation of all time series representing the system. | maxSD |
| | min/max autocorrelation factor (MAF) dominant eigenvalue | Following MAF dimension reduction of all representative time series, smallest scalar | eigenMAF |

| | | | |
|---|---|---|---|
| | | of the resulting eigenvectors. | |
| | first MAF (MAF1) autocorrelation at lag-1 | The autocorrelation of the MAF axis that yields the strongest autocorrelation. | mafAR |
| | MAF1 variance | The standard deviation of the MAF axis that yields the strongest autocorrelation. | mafSD |
| | first principal component (PC1) autocorrelation at lag-1 | Following principal component analysis of all representative time series, the autocorrelation of the principal component which explains the greatest variance. | pcaAR |
| | PC1 variance | The standard deviation of the first principal component. | pcaSD |
| | dominant eigenvalue of the covariance matrix | From the covariance matrix of all representative time series, the largest eigenvalue is informative. | eigenCOV |
| | maximum covariance | The strongest covariance between all representative time series. | maxCOV |
| | mutual information | The degree of information gained from one time series on the state of another. Is averaged across all pairwise time series comparisons. | mutINFO |
| Machine learning | EWSNet scaled weights | Calls the scaled forms of the EWSNet model weights (scaled between 1-2) and conceptually should be robust regardless of the data’s magnitude. | scaled |
| | EWSNet unscaled weights | Calls the unscaled forms of the EWSNet model weights. | unscaled |

**Table S3.** Coefficient estimates for influence of each data pre-processing technique on rolling window univariate early warning signal classification ability relative to assessments made on the raw data.

| Pre-processing combination<br><br>(detrending method – deseasoning method) | Estimated improvement (median) | Lower 95% credible interval | Upper 95% credible interval | Rhat | Effective sample size |
|---|---|---|---|---|---|
| linear-none | 0.021 | -0.293 | 0.336 | 1 | 4591.67 |
| loess-none | -0.034 | -0.335 | 0.273 | 1 | 4999.17 |
| gaussian-none | -0.047 | -0.363 | 0.278 | 1 | 5235.38 |
| none-average | 0.021 | -0.291 | 0.336 | 1 | 5048.17 |
| none-decomposition | 0.052 | -0.262 | 0.355 | 1 | 4600.02 |
| none-stl | -0.021 | -0.33 | 0.292 | 1 | 5131.11 |
| linear-average | 0.006 | -0.3 | 0.323 | 1 | 5248.49 |
| loess-average | 0.037 | -0.28 | 0.352 | 1 | 5081.21 |
| gaussian-average | 0.023 | -0.293 | 0.341 | 1 | 5033.33 |
| linear-decomposition | 0.036 | -0.284 | 0.352 | 1 | 5097.07 |
| loess-decomposition | 0.011 | -0.294 | 0.33 | 1 | 4660.88 |
| gaussian-decomposition | 0.005 | -0.298 | 0.33 | 1 | 5274.77 |
| linear-stl | 0.079 | -0.23 | 0.402 | 1 | 4755.66 |
| loess-stl | 0.018 | -0.283 | 0.336 | 1 | 4600.65 |
| gaussian-stl | 0.034 | -0.276 | 0.345 | 1 | 4712.69 |

**Table S4.** Coefficient estimates for influence of each data pre-processing technique on expanding window univariate early warning signal classification ability relative to assessments made on the raw data.

| **Pre-processing combination**<br><br>**(detrending method – deseasoning method)** | **Estimated improvement (median)** | **Lower 95% credible interval** | **Upper 95% credible interval** | **Rhat** | **Effective sample size** |
|---|---|---|---|---|---|
| linear-none | 0.009 | -0.204 | 0.225 | 1 | 4749.32 |
| loess-none | -0.083 | -0.291 | 0.127 | 1 | 4966.58 |
| gaussian-none | -0.164 | -0.371 | 0.046 | 1 | 5276.24 |
| none-average | -0.1 | -0.31 | 0.105 | 1 | 4989.79 |
| none-decomposition | 0.209 | -0.002 | 0.425 | 1 | 4398.79 |
| none-stl | -0.008 | -0.22 | 0.203 | 1 | 4817.7 |
| linear-average | -0.086 | -0.297 | 0.125 | 1 | 4388.72 |
| loess-average | -0.212 | -0.415 | -0.006 | 1 | 5299.05 |
| gaussian-average | -0.283 | -0.489 | -0.078 | 1 | 5029.48 |
| linear-decomposition | 0.184 | -0.032 | 0.392 | 1 | 5075.47 |
| loess-decomposition | -0.072 | -0.281 | 0.139 | 1 | 4682.55 |
| gaussian-decomposition | -0.089 | -0.295 | 0.119 | 1 | 4677.83 |
| linear-stl | -0.082 | -0.293 | 0.121 | 1 | 4777.39 |
| loess-stl | -0.153 | -0.358 | 0.05 | 1 | 4730.88 |
| gaussian-stl | -0.232 | -0.437 | -0.032 | 1 | 4959.17 |

**Table S5.** Coefficient estimates for influence of each data pre-processing technique on rolling window multivariate early warning signal classification ability relative to assessments made on the raw data.

| **Pre-processing combination**<br><br>**(detrending method – deseasoning method)** | **Estimated improvement (median)** | **Lower 95% credible interval** | **Upper 95% credible interval** | **Rhat** | **Effective sample size** |
|---|---|---|---|---|---|
| linear-none | 0.116 | -0.422 | 0.664 | 1 | 5306.62 |
| loess-none | 0.161 | -0.376 | 0.706 | 1 | 5736.47 |
| gaussian-none | 0.208 | -0.347 | 0.756 | 1 | 5833.45 |
| none-average | -0.39 | -0.947 | 0.154 | 1 | 5367.25 |
| none-decomposition | -0.299 | -0.848 | 0.247 | 1 | 5748.78 |
| none-stl | -0.201 | -0.744 | 0.341 | 1 | 5425.41 |
| linear-average | -0.206 | -0.743 | 0.336 | 1 | 5552.16 |
| loess-average | -0.069 | -0.606 | 0.466 | 1 | 5366.18 |
| gaussian-average | -0.071 | -0.595 | 0.481 | 1 | 5498.8 |
| linear-decomposition | -0.067 | -0.612 | 0.465 | 1 | 5651.47 |
| loess-decomposition | -0.067 | -0.619 | 0.458 | 1 | 5725.4 |
| gaussian-decomposition | 0.162 | -0.376 | 0.7 | 1 | 5243.71 |
| linear-stl | -0.022 | -0.564 | 0.513 | 1 | 5567.68 |
| loess-stl | 0.07 | -0.473 | 0.599 | 1 | 5771.72 |
| gaussian-stl | -0.023 | -0.572 | 0.528 | 1 | 5713.59 |

**Table S6.** Coefficient estimates for influence of each data pre-processing technique on expanding window multivariate early warning signal classification ability relative to assessments made on the raw data.

| **Pre-processing combination**<br><br>**(detrending method – deseasoning method)** | **Estimated improvement (median)** | **Lower 95% credible interval** | **Upper 95% credible interval** | **Rhat** | **Effective sample size** |
|---|---|---|---|---|---|
| linear-none | -0.072 | -0.637 | 0.5 | 1 | 5266.96 |
| loess-none | -0.022 | -0.579 | 0.588 | 1 | 5511.38 |
| gaussian-none | 0.139 | -0.422 | 0.726 | 1 | 4846.59 |
| none-average | -0.118 | -0.691 | 0.454 | 1 | 5351.54 |
| none-decomposition | 0.149 | -0.437 | 0.718 | 1 | 5092.31 |
| none-stl | -0.124 | -0.674 | 0.447 | 1 | 5009.86 |
| linear-average | -0.124 | -0.679 | 0.447 | 1 | 5641.68 |
| loess-average | -0.124 | -0.695 | 0.446 | 1 | 5351.4 |
| gaussian-average | 0.242 | -0.329 | 0.83 | 1 | 4943.64 |
| linear-decomposition | -0.333 | -0.895 | 0.238 | 1 | 5451.45 |
| loess-decomposition | -0.486 | -1.047 | 0.074 | 1 | 5084.2 |
| gaussian-decomposition | 0.038 | -0.533 | 0.609 | 1 | 5423.48 |
| linear-stl | -0.076 | -0.65 | 0.5 | 1 | 4735.91 |
| loess-stl | -0.278 | -0.846 | 0.297 | 1 | 5487.01 |
| gaussian-stl | 0.135 | -0.42 | 0.713 | 1 | 5093.32 |

**Table S7.** Coefficient estimates for influence of each data pre-processing technique on machine learning univariate early warning signal classification ability relative to assessments made on the raw data.

| **Pre-processing combination**<br><br>**(detrending method – deseasoning method)** | **Estimated improvement (median)** | **Lower 95% credible interval** | **Upper 95% credible interval** | **Rhat** | **Effective sample size** |
|---|---|---|---|---|---|
| linear-none | 0.08 | -0.309 | 0.46 | 1 | 5530.66 |
| loess-none | 0.181 | -0.199 | 0.571 | 1 | 5156.51 |
| gaussian-none | 0.223 | -0.16 | 0.605 | 1 | 5147.1 |
| none-average | -0.028 | -0.415 | 0.365 | 1 | 5117.16 |
| none-decomposition | -0.138 | -0.515 | 0.254 | 1 | 4831.59 |
| none-stl | -0.087 | -0.462 | 0.287 | 1 | 4949.26 |
| linear-average | 0.097 | -0.288 | 0.478 | 1 | 5028.33 |
| loess-average | 0.077 | -0.311 | 0.468 | 1 | 5284.37 |
| gaussian-average | 0.184 | -0.197 | 0.571 | 1 | 5401.19 |
| linear-decomposition | 0.099 | -0.28 | 0.473 | 1 | 5236.98 |
| loess-decomposition | 0.02 | -0.371 | 0.398 | 1 | 5105.53 |
| gaussian-decomposition | 0.101 | -0.277 | 0.478 | 1 | 4991.41 |
| linear-stl | -0.202 | -0.593 | 0.176 | 1 | 4931.48 |
| loess-stl | -0.008 | -0.389 | 0.37 | 1 | 5021.52 |
| gaussian-stl | -0.01 | -0.392 | 0.378 | 1 | 4896.23 |

**Table S8.** Raw coefficient estimates for the influence of each early warning signal computational approach on correct classification of the monthly lake plankton dataset.

| Computation method | Estimated improvement (median) | Lower 95% credible interval | Upper 95% credible interval | Rhat | Effective sample size |
|---|---|---|---|---|---|
| univariate_rolling | 0.27 | 0.009 | 0.495 | 1 | 6637.18 |
| univariate_expanding | 0.532 | 0.277 | 0.741 | 1 | 5965.87 |
| EWSNet | -0.101 | -0.376 | 0.148 | 1 | 6669.06 |
| multivariate_rolling | 0.132 | -0.199 | 0.45 | 1 | 6579.43 |
| multivariate_expanding | 0.592 | 0.262 | 0.929 | 1 | 6709.26 |

**Table S9.** Raw coefficient estimates for the influence of each early warning signal computational approach on correct classification of the yearly lake plankton dataset.

| Computation method | Estimated improvement (median) | Lower 95% credible interval | Upper 95% credible interval | Rhat | Effective sample size |
|---|---|---|---|---|---|
| univariate_rolling | -0.759 | -1.53 | -0.055 | 1 | 6738.84 |
| univariate_expanding | 2.171 | 1.384 | 2.914 | 1 | 6677.44 |
| EWSNet | -0.243 | -1.013 | 0.457 | 1 | 6605.16 |
| multivariate_rolling | -0.493 | -1.285 | 0.244 | 1 | 6744.16 |
| multivariate_expanding | 1.056 | 0.217 | 1.847 | 1 | 6596.97 |

**Table S10.** Raw coefficient estimates for the influence of each early warning signal indicator on the correct classification of transitioning monthly lake plankton data.

| Early warning signal indicator | Estimated improvement (median) | Lower 95% credible interval | Upper 95% credible interval | Rhat | Effective sample size |
|---|---|---|---|---|---|
| ar1 + SD + skew_expanding | -0.112 | -1.236 | 1.032 | 1 | 6510.13 |
| ar1 + SD_expanding | 0.095 | -1.027 | 1.198 | 1 | 6742.26 |
| ar1 + skew_expanding | -0.315 | -1.432 | 0.81 | 1 | 6972.55 |
| ar1_expanding | -0.292 | -1.423 | 0.79 | 1 | 6721.85 |
| ar1_rolling | 0.226 | -0.785 | 1.21 | 1 | 6986.89 |
| eigenCOV_expanding | 0.075 | -1.577 | 1.768 | 1 | 7329.82 |
| eigenCOV_rolling | 1.266 | -0.449 | 3.126 | 1 | 6284.91 |
| eigenMAF_expanding | 0.079 | -1.633 | 1.772 | 1 | 6875.22 |
| eigenMAF_rolling | 0.532 | -1.125 | 2.298 | 1 | 7437.93 |
| mafAR_expanding | 0.08 | -1.591 | 1.827 | 1 | 7198.22 |
| mafAR_rolling | -0.125 | -1.783 | 1.55 | 1 | 7010.8 |
| mafSD_expanding | -0.622 | -2.414 | 1.096 | 1 | 6952.79 |
| mafSD_rolling | 1.261 | -0.413 | 3.146 | 1 | 7204.08 |
| maxAR_expanding | 0.787 | -0.917 | 2.569 | 1 | 7018.18 |
| maxAR_rolling | -0.786 | -2.486 | 0.82 | 1 | 7189.31 |
| maxCOV_expanding | 0.095 | -1.552 | 1.8 | 1 | 7274.56 |
| maxCOV_rolling | 0.53 | -1.148 | 2.262 | 1 | 6958.78 |
| maxSD_expanding | -0.629 | -2.415 | 1.058 | 1 | 6894.74 |
| maxSD_rolling | -1.523 | -3.293 | 0.154 | 1 | 6933.54 |
| meanAR_expanding | 0.09 | -1.588 | 1.765 | 1 | 7127.01 |
| meanAR_rolling | 0.529 | -1.161 | 2.284 | 1 | 7114.28 |
| meanSD_expanding | 0.068 | -1.631 | 1.77 | 1 | 6486.73 |
| meanSD_rolling | -0.798 | -2.525 | 0.835 | 1 | 7051.88 |
| mutINFO_expanding | -1.372 | -3.259 | 0.291 | 1 | 7099.94 |
| mutINFO_rolling | -0.134 | -1.782 | 1.536 | 1 | 7396.25 |
| pcaAR_expanding | 0.09 | -1.6 | 1.812 | 1 | 7303.65 |
| pcaAR_rolling | -0.131 | -1.735 | 1.528 | 1 | 7120.27 |
| pcaSD_expanding | 0.085 | -1.642 | 1.739 | 1 | 6874.56 |
| pcaSD_rolling | 1.28 | -0.415 | 3.162 | 1 | 7137.58 |
| scaled_ML | -1.919 | -3.17 | -0.805 | 1 | 6947.11 |
| SD + skew_expanding | -0.112 | -1.207 | 0.988 | 1 | 6994.47 |
| SD_expanding | 0.307 | -0.781 | 1.392 | 1 | 7065.92 |
| SD_rolling | -0.339 | -1.339 | 0.617 | 1 | 6306.79 |
| skew_expanding | -0.505 | -1.587 | 0.596 | 1 | 6883.78 |
| skew_rolling | -0.225 | -1.232 | 0.764 | 1 | 7068.66 |
| unscaled_ML | 2.559 | 1.309 | 3.953 | 1 | 7170.79 |

**Table S11.** Raw coefficient estimates for the influence of each early warning signal indicator on the correct classification of transitioning yearly lake plankton data.

| Early warning signal indicator | Estimated improvement (median) | Lower 95% credible interval | Upper 95% credible interval | Rhat | Effective sample size |
|---|---|---|---|---|---|
| ar1 + SD_expanding | 0.572 | -1.437 | 2.439 | 1 | 7047.26 |
| ar1 + skew_expanding | -0.408 | -2.538 | 1.608 | 1 | 6992.48 |
| ar1_expanding | -0.406 | -2.524 | 1.553 | 1 | 7670.5 |
| ar1_rolling | 0.172 | -1.199 | 1.515 | 1 | 7068.68 |
| eigenCOV_expanding | -0.25 | -2.525 | 1.828 | 1 | 7251.68 |
| eigenCOV_rolling | 0.923 | -0.795 | 2.643 | 1 | 7166.63 |
| eigenMAF_expanding | -0.237 | -2.397 | 1.83 | 1 | 7260.28 |
| eigenMAF_rolling | 0.923 | -0.816 | 2.659 | 1 | 7165.82 |
| mafAR_expanding | -0.24 | -2.464 | 1.875 | 1 | 6856.17 |
| mafAR_rolling | 0.232 | -1.453 | 1.939 | 1 | 6901.05 |
| mafSD_expanding | -0.231 | -2.434 | 1.857 | 1 | 6818.39 |
| mafSD_rolling | 0.249 | -1.481 | 1.937 | 1 | 7209.95 |
| maxAR_expanding | -0.251 | -2.459 | 1.867 | 1 | 7297.64 |
| maxAR_rolling | -0.477 | -2.209 | 1.217 | 1 | 7387.4 |
| maxCOV_expanding | -0.253 | -2.465 | 1.789 | 1 | 7216.45 |
| maxCOV_rolling | 0.905 | -0.749 | 2.649 | 1 | 7230.45 |
| maxSD_expanding | -0.222 | -2.512 | 1.889 | 1 | 7323.75 |
| maxSD_rolling | -0.454 | -2.261 | 1.258 | 1 | 7075.98 |
| meanAR_expanding | -0.234 | -2.449 | 1.831 | 1 | 7054.79 |
| meanAR_rolling | 0.221 | -1.521 | 1.904 | 1 | 7148.85 |
| meanSD_expanding | -0.268 | -2.454 | 1.859 | 1 | 7178.53 |
| meanSD_rolling | -0.462 | -2.286 | 1.186 | 1 | 7112 |
| mutINFO_expanding | -0.248 | -2.47 | 1.872 | 1 | 7236.54 |
| mutINFO_rolling | -1.211 | -3.128 | 0.496 | 1 | 7054.34 |
| pcaAR_expanding | -0.272 | -2.432 | 1.851 | 1 | 7251.23 |
| pcaAR_rolling | -0.473 | -2.242 | 1.243 | 1 | 7197.75 |
| pcaSD_expanding | -0.269 | -2.399 | 1.887 | 1 | 6907.13 |
| pcaSD_rolling | 0.912 | -0.82 | 2.692 | 1 | 7124.44 |
| scaled_ML | -2.558 | -4.38 | -0.882 | 1 | 7169.49 |
| SD + skew_expanding | -0.402 | -2.517 | 1.555 | 1 | 7211.07 |
| SD_expanding | -0.378 | -2.531 | 1.491 | 1 | 7103.37 |
| SD_rolling | 0.404 | -0.952 | 1.709 | 1 | 7002.61 |
| skew_expanding | -0.393 | -2.513 | 1.587 | 1 | 7342.03 |
| skew_rolling | 0.513 | -0.874 | 1.852 | 1 | 7251.06 |
| unscaled_ML | 2.092 | 0.516 | 3.682 | 1 | 6851.29 |
| ar1 + SD_expanding | 0.572 | -1.437 | 2.439 | 1 | 7047.26 |

**Table S12.** Raw coefficient estimates for the influence of each early warning signal indicator on the correct classification of non-transitioning monthly lake plankton data.

| Early warning signal indicator | Estimated improvement (median) | Lower 95% credible interval | Upper 95% credible interval | Rhat | Effective sample size |
|---|---|---|---|---|---|
| ar1 + SD_expanding | 0.369 | 0.027 | 0.69 | 1 | 6934.22 |
| ar1 + skew_expanding | 1.207 | 0.841 | 1.569 | 1 | 6905.81 |
| ar1_expanding | 0.742 | 0.389 | 1.07 | 1 | 6505.07 |
| ar1_rolling | 0.285 | -0.044 | 0.599 | 1 | 7070.57 |
| eigenCOV_expanding | 1.366 | 0.282 | 2.644 | 1 | 6777.09 |
| eigenCOV_rolling | -0.521 | -1.536 | 0.468 | 1 | 7259.13 |
| eigenMAF_expanding | 0.214 | -0.739 | 1.22 | 1 | 7288.79 |
| eigenMAF_rolling | -0.018 | -1.015 | 0.971 | 1 | 6803.98 |
| mafAR_expanding | 0.219 | -0.769 | 1.233 | 1 | 6737.03 |
| mafAR_rolling | -0.017 | -1.005 | 0.978 | 1 | 6839.93 |
| mafSD_expanding | -0.271 | -1.298 | 0.712 | 1 | 7612.75 |
| mafSD_rolling | -0.511 | -1.581 | 0.464 | 1 | 6549.93 |
| maxAR_expanding | 0.23 | -0.744 | 1.222 | 1 | 7107.68 |
| maxAR_rolling | -0.017 | -1.006 | 0.972 | 1 | 7329.67 |
| maxCOV_expanding | 0.482 | -0.547 | 1.551 | 1 | 7196.94 |
| maxCOV_rolling | -0.254 | -1.298 | 0.715 | 1 | 7055.02 |
| maxSD_expanding | 0.751 | -0.265 | 1.846 | 1 | 7310.43 |
| maxSD_rolling | 0.761 | -0.238 | 1.839 | 1 | 7083.14 |
| meanAR_expanding | 1.72 | 0.598 | 3.125 | 1 | 6763.13 |
| meanAR_rolling | -0.256 | -1.271 | 0.719 | 1 | 7091.02 |
| meanSD_expanding | 0.21 | -0.769 | 1.236 | 1 | 7328.28 |
| meanSD_rolling | 1.372 | 0.281 | 2.661 | 1 | 6921.84 |
| mutINFO_expanding | 2.206 | 0.944 | 3.711 | 1 | 7279.62 |
| mutINFO_rolling | 0.241 | -0.727 | 1.271 | 1 | 7432.17 |
| pcaAR_expanding | 1.04 | -0.006 | 2.206 | 1 | 7510.32 |
| pcaAR_rolling | 0.502 | -0.483 | 1.536 | 1 | 7403.45 |
| pcaSD_expanding | 1.382 | 0.284 | 2.647 | 1 | 6788.08 |
| pcaSD_rolling | -0.529 | -1.586 | 0.467 | 1 | 7017.65 |
| scaled_ML | 1.174 | 0.804 | 1.529 | 1 | 7088.68 |
| SD + skew_expanding | 0.659 | 0.312 | 0.985 | 1 | 6189.2 |
| SD_expanding | 0.04 | -0.294 | 0.342 | 1 | 7019.35 |
| SD_rolling | 0.608 | 0.257 | 0.935 | 1 | 6988.25 |
| skew_expanding | 1.148 | 0.793 | 1.502 | 1 | 6134.02 |
| skew_rolling | 0.302 | -0.025 | 0.613 | 1 | 6604.9 |
| unscaled_ML | -1.419 | -1.829 | -1.05 | 1 | 7145.85 |
| ar1 + SD_expanding | 0.369 | 0.027 | 0.69 | 1 | 6934.22 |

**Table S13.** Raw coefficient estimates for the influence of each early warning signal indicator on the correct classification of non-transitioning yearly lake plankton data.

| Early warning signal indicator | Estimated improvement (median) | Lower 95% credible interval | Upper 95% credible interval | Rhat | Effective sample size |
|---|---|---|---|---|---|
| ar1 + SD_expanding | 0.647 | -0.525 | 1.866 | 1 | 7000.68 |
| ar1 + skew_expanding | 0.259 | -0.84 | 1.43 | 1 | 7033.54 |
| ar1_expanding | 1.144 | -0.115 | 2.511 | 1 | 7037.47 |
| ar1_rolling | -1.157 | -2.277 | 0.034 | 1 | 6616.09 |
| eigenCOV_expanding | 0.514 | -1.344 | 2.598 | 1 | 7422.24 |
| eigenCOV_rolling | -0.79 | -1.977 | 0.417 | 1 | 6883.87 |
| eigenMAF_expanding | 0.498 | -1.418 | 2.618 | 1 | 7185.15 |
| eigenMAF_rolling | -0.217 | -1.421 | 1.022 | 1 | 7113.81 |
| mafAR_expanding | 0.494 | -1.395 | 2.66 | 1 | 7127.44 |
| mafAR_rolling | -0.515 | -1.702 | 0.713 | 1 | 7089.76 |
| mafSD_expanding | 0.489 | -1.377 | 2.637 | 1 | 7230.03 |
| mafSD_rolling | -1.368 | -2.64 | -0.177 | 1 | 6849.81 |
| maxAR_expanding | 0.481 | -1.407 | 2.627 | 1 | 6749.45 |
| maxAR_rolling | 0.711 | -0.551 | 2.049 | 1 | 7254.52 |
| maxCOV_expanding | 0.519 | -1.385 | 2.657 | 1 | 7150.11 |
| maxCOV_rolling | -0.783 | -2.01 | 0.414 | 1 | 6964.69 |
| maxSD_expanding | 0.518 | -1.369 | 2.642 | 1 | 7085 |
| maxSD_rolling | 1.508 | 0.136 | 3.007 | 1 | 6954.06 |
| meanAR_expanding | 0.524 | -1.393 | 2.584 | 1 | 6970.53 |
| meanAR_rolling | 0.08 | -1.138 | 1.333 | 1 | 7069.7 |
| meanSD_expanding | 0.481 | -1.356 | 2.616 | 1 | 7223.43 |
| meanSD_rolling | 1.082 | -0.228 | 2.504 | 1 | 7023.5 |
| mutINFO_expanding | 0.522 | -1.397 | 2.613 | 1 | 6931.83 |
| mutINFO_rolling | -2.859 | -4.401 | -1.474 | 1 | 7039.7 |
| pcaAR_expanding | 0.508 | -1.4 | 2.652 | 1 | 7238.16 |
| pcaAR_rolling | 0.071 | -1.142 | 1.33 | 1 | 7057.4 |
| pcaSD_expanding | 0.504 | -1.426 | 2.579 | 1 | 6907.75 |
| pcaSD_rolling | -0.784 | -1.992 | 0.39 | 1 | 6316.35 |
| scaled_ML | 3.639 | 2.156 | 5.093 | 1 | 6651.81 |
| SD + skew_expanding | 0.441 | -0.684 | 1.63 | 1 | 6952.17 |
| SD_expanding | 0.645 | -0.551 | 1.897 | 1 | 6876.36 |
| SD_rolling | -0.565 | -1.684 | 0.643 | 1 | 6633.89 |
| skew_expanding | 1.5 | 0.186 | 3.01 | 1 | 6899.53 |
| skew_rolling | -0.915 | -2.021 | 0.283 | 1 | 6799.71 |
| unscaled_ML | -2.937 | -4.221 | -1.683 | 1 | 7267.15 |
| ar1 + SD_expanding | 0.647 | -0.525 | 1.866 | 1 | 7000.68 |
